# Glymphatic Optimal Mass Transport with Lagrangian Workflow Reveals Advective and Diffusion Driven Solute Transport

**DOI:** 10.1101/765370

**Authors:** Sunil Koundal, Rena Elkin, Saad Nadeem, Yuechuan Xue, Stefan Constantinou, Simon Sanggaard, Xiaodan Liu, Brittany Monte, Feng Xu, William Van Nostrand, Maiken Nedergaard, Hedok Lee, Joanna Wardlaw, Helene Benveniste, Allen Tannenbaum

**Author notes:** These authors contributed equally to this work (Sunil Koundal, Rena Elkin). Co-Senior Authors (Helene Benveniste, Allen Tannenbaum). **Author contributions:** AT and HB conceived the study; AT and RE developed the rOMT algorithm and Lagrangian framework analysis; RE performed all rOMT processing; SN provided key suggestions for processing the Lagrangian analysis; HB performed all rOMT post-processing and designed figures with contributions from RE; HB performed kinetic analysis; SK, SS and XL performed all glymphatics experiments. SK executed all morphometric analysis including brain atlas segmentation; HL designed all pulse-sequences and other hardware for the MRI experiments and computational pipeline for the volumetric analysis. SS, FX, WVN performed the immunohistochemistry. SC performed qPCR analysis. YX performed the quantitative AQP4 analysis. HB, RE and AT wrote the manuscript. JW advised on the SHRSP animal model and cerebral small vessel disease. MN advised on the glymphatic system. All authors posed scientific questions, read and revised the manuscript. All authors edited and reviewed the paper.

## Abstract

The presence of advection in neuropil is contested and solute transport is claimed to occur by diffusion only. To address this controversy, we implemented a regularized version of the optimal mass transport (rOMT) problem, wherein the advection/diffusion equation is the only *a priori* assumption required. rOMT analysis with a Lagrangian perspective of glymphatic system (GS) transport revealed that solute speed was faster in cerebrospinal fluid (CSF) compared to grey and white matter. rOMT analysis also demonstrated 2-fold differences in regional particle speed within the brain parenchyma. Collectively, these results imply that advective transport dominates in CSF while diffusion and advection both contribute to transport in parenchyma. In rats with chronic hypertension, solute transport in perivascular spaces (PVS) and PVS-to-tissue transfer was slower compared to normotension. Thus, the analytical framework of rOMT provides novel insights in local variation and dynamics of GS transport that may have implications for neurodegenerative diseases.

## Introduction

The glymphatic system (GS) is described as a perivascular transit passageway for cerebrospinal fluid (CSF) for exchange with interstitial fluid (ISF), thereby facilitating waste drainage from the brain^1, 2^. Investigations of GS function have escalated given its important role in Aβ^1^ and tau^3^ drainage from brain and the inferred implication for neurodegeneration including Alzheimer’s disease^2, 4–7^. The glymphatic system (GS) hypothesis states that *advective* driven CSF influx from the perivascular spaces (PVS) into brain parenchyma clears solutes^1, 8^. Advective streams in parenchyma would speed up solute transport^9–11^. This concept is contested with the argument that advection does not occur in the neuropil and that solute transport occurs by diffusion only^12–18^. No single study has unequivocally determined which mode of solute transport is prevailing inside the brain proper. This gap in knowledge has significantly impeded exploration of the GS and has raised questions regarding its importance for brain health, disease, and its potential as a therapeutic target against neurodegeneration.

The GS is comprised of PVS, which connect with the ISF space surrounding all cells in parenchyma^1^. Specifically, the PVS connect with ISF via the aquaporin 4 (AQP4) water channels on astrocytic end-feet and through the small gaps between the overlapping astrocytic end-foot processes^1^. Solute transport in the PVS along pial arteries has been shown to be an advective-driven (bulk flow) process^1, 19–21^. Recently, various computational analyses and simulations have been performed to better understand the driving mechanisms of transport across PVS boundaries and through ISF space^22–24^. Conservation principles are typically used to derive these simulations, a popular one being the porous media model^25–27^. Ray et al.^26^ proposed adding an advection term of this form to the (diffusive) porous media model and their simulations indicated the presence of advective transport in the ISF. Simulations are very effective for distinguishing between advection and diffusion, but when applied *in vivo*, inferable insight becomes limited. The key reason for this problem, lies in the fact that modeling in live brain requires accurate delineation of anatomical boundaries and knowledge of various kinematic parameters many of which are unknown.

To the best of our knowledge, no current method can provide local dynamic analysis or inform on GS transport modes across compartments in the live brain. Therefore, we now introduce novel computational methods to improve the interpretation of GS transport studies. Specifically, we address two overarching and unresolved questions: (1) Can we obtain *voxel-level* information on solute transport across all tissue compartments circumventing the problem of varying GS transport kinetics and without the need of specifying all of the material properties for simulations? (2) Can we infer on the existence of the two types of transport (*diffusion and advection*) in the different GS compartments? To address these questions, we propose implementing techniques from the theory of optimal mass transport (OMT)^28–30^. The OMT problem is centuries old and has impacted many topics of research in the physical sciences, economics, computer science and image processing^28, 31–33^. Given initial and final mass distributions, OMT admits a powerful dynamic computational fluid formulation that gives an optimal interpolation path of minimal energy among all possible interpolations that preserve mass^34^. In this way, OMT can be formulated as a variational problem based on the principle of least action. A particularly attractive feature of OMT is that one obtains unique solutions to the model, not by imposing smoothness, but rather by seeking flows that minimize the expended transport energy. In the classical formulation of OMT, the continuity equation only involves advection. Considering that diffusion is assumed to be always occurring in the brain, we have added a diffusion term in the work presented here. We refer to our modified OMT formulation with the advection/diffusion equation as the *regularized OMT problem* (rOMT). We utilized an associated Lagrangian formulation of rOMT for the construction of ‘pathlines’ to effectively extract and visualize GS transport flows over a predetermined set of time frames in one comprehensive figure. Time-varying particle (a.k.a. solute) attributes associated with the pathlines, such as ‘speed’ were also computed. As GS dysfunction is reported in neurodegeneration particularly where vascular dysfunction may be involved^7, 35, 36^, we applied our novel rOMT framework to characterize modes of solute transport in an animal model of cerebral small vessel disease.

In normal brains, rOMT analysis revealed that both advection and diffusion terms were required for pathlines and trajectory speed in the modeled images to resemble GS transport patterns in live brain. Moreover, particle speed varied strikingly across tissue compartments with faster solute passage in CSF when compared to parenchyma. Within the brain parenchyma, 2-fold regional differences in solute speed were also documented. Collectively, these novel results in live rat brain demonstrate that advective transport is dominating in the non-cellular CSF spaces and that slower diffusion contributes to GS transport as the solute is transferred from the PVS into ISF. Interestingly, in chronic hypertension, PVS-CSF flow and solute transport across the PVS into parenchyma were impaired. We believe that our new rOMT Lagrangian framework analysis will prove informative for characterizing local variation in GS transport in neurogenerative states that would otherwise remain controversial.

## Results

### Kinetic analysis of GS transport in normal and hypertensive rats

We first characterized GS transport in normotensive Wistar Kyoto (WKY) and spontaneous hypertensive stroke prone (SHRSP) rats using kinetic modeling^8, 37, 38^. To measure GS transport, we used a dynamic contrast enhanced magnetic resonance image (DCE-MRI) technique by infusion of 559 Da sized gadoteric acid (Gd-DOTA, 20 μL of 1:37 Gd-DOTA diluted with sterile water at a rate of 1.5 μL/min) into the CSF via the cisterna magna^39^. DCE-MRIs covering the entire brain were acquired every 5-min for 2.5 hrs on a 9.4T MRI at a voxel resolution 0.30×0.30×0.30 mm^3^. In normal rats, the tracer was observed in the subarachnoid CSF space along the large basilar artery, circle of Willis and olfactory artery within 10-15 min and later emerged along branches of the posterior cerebral arteries which penetrate into the hippocampus and thalamus (**Figs. 1a, b**). Voxel-based morphometric segmentation of each brain into CSF, grey matter (GM) and white matter (WM) tissue masks derived from anatomical proton-weighted MRIs of the two rat strains (see methods) were used to calculate influx and efflux rate constants as well as tracer distribution volumes across brain compartments. A known feature of the spontaneously hypertensive rat (SHR) strain used in previous GS studies^37, 40^ is innate hydrocephalus^41, 42^, which can affect CSF fluid dynamics long-term^43^. However, no major morphometric differences were observed between WKY and SHRSP rats, and hydrocephalus was not present in SHRSP rats (**Supplementary Fig. 1**).

**Figure 1:**
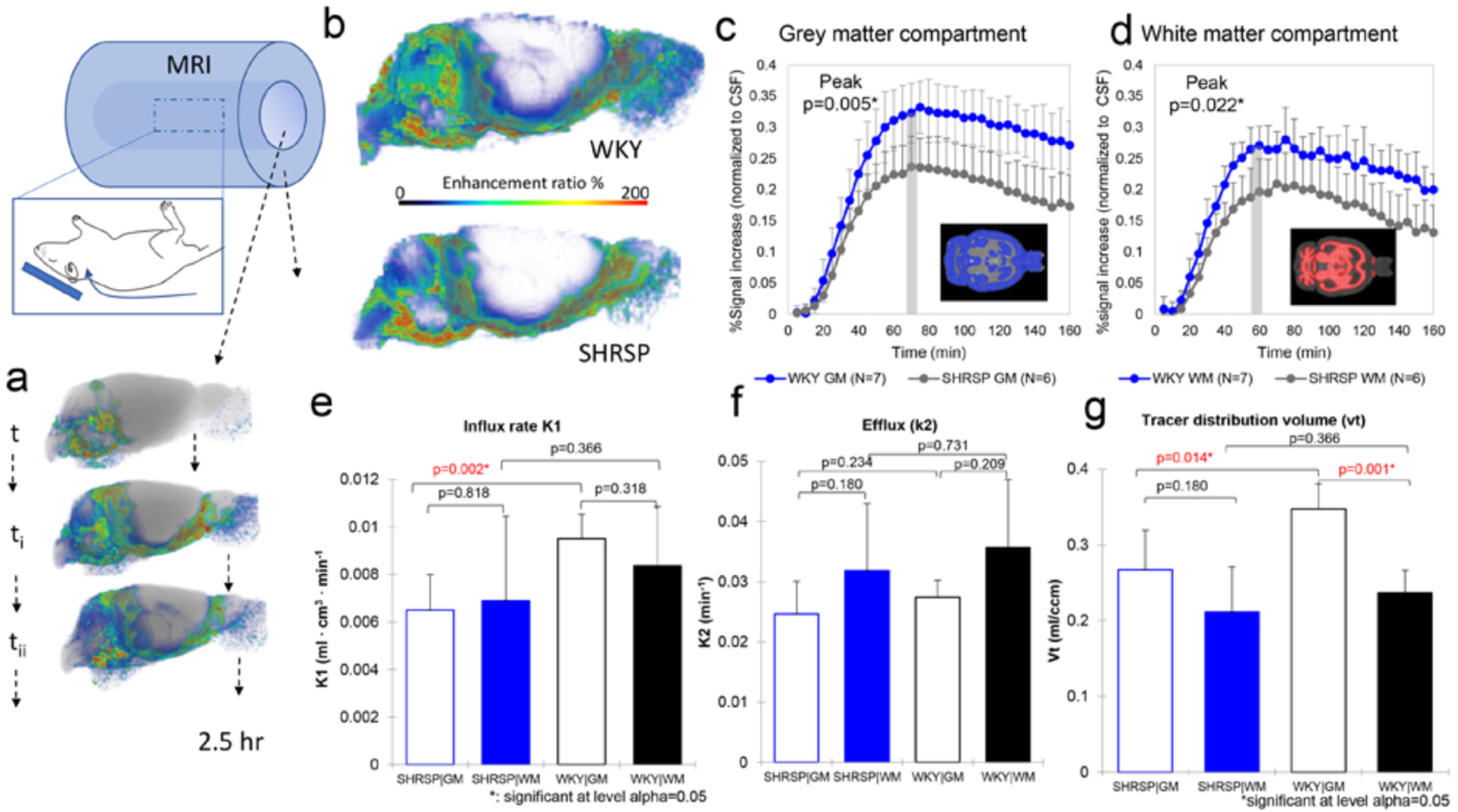
Kinetic modeling of GS transport in normotensive and hypertensive rats. **(a)** Glymphatic transport was visualized and measured in live rats through administration of Gd-DOTA into CSF via the cisterna magna using dynamic contrast enhanced magnetic resonance imaging (DCE-MRI). 3D DCE-MRIs were acquired every 5 min to track Gd-DOTA transport (measured as % signal changes from baseline) into CSF, white matter (WM) and grey matter (GM) compartments over 2.5 hrs. **(b)** Volume-rendered color-coded maps of Gd-DOTA induced signal change over 110 min in a WKY rat (top) and SHRSP rat (bottom). It is clear that Gd-DOTA uptake is reduced in the SHRSP rat when compared to WKY. (**c)** Mean signal changes (normalized to signal in CSF compartment) captured in the GM compartment of WKY (blue circles) and SHRSP (gray circles) rats over the 160 min experimental time window. Insert: grey matter mask is shown in blue overlaid on the anatomical MRI. Peak magnitude in GM at ~70 min of circulation was significantly decreased in SHRSP when compared to WKY (p=0.022). (**d**) Mean signal changes (normalized to signal in CSF compartment) in WM of WKY (blue circles) and SHRSP (gray circles) over the 160 min experimental time window. The insert shows the WM mask in red overlaid on the anatomical template. Peak magnitude in WM at ~60 min of circulation was significantly decreased in SHRSP when compared to WKY (p=0.022). **(e)** Mean K1 influx rate calculated from 1-compartment kinetic modeling is significantly reduced in GM but not WM of SHRSP compared to WKY rats. (**f)** Mean efflux rates were similar across GM and WM compartments and no differences were observed across strains. (**g)** Distribution volumes of Gd-DOTA were significantly higher in GM and WM of WKY when compared to SHRSP rats. Data are mean ± SD. Mann-Whitney test, *significant at level alpha=0.05.

Globally, brain tissue uptake of Gd-DOTA was strikingly different between the two strains (**Fig. 1b**) with reduced uptake in both GM and WM compartments of SHRSP when compared to WKY rats (**Figs. 1c, d**). Peak signal magnitude in GM and WM was significantly reduced in SHRSP when compared to WKY rats **(Fig. 1c, d)**. Kinetic one-compartment modelling with the input given as the MRI measure of time-varying Gd-DOTA-induced signal levels in CSF, GM and WM compartments (**Figs. 1c, d**, **inserts**)were used to calculate the influx coefficient K1 and efflux rate constant k2 as well as the distribution volume of the tracer V(t)^37^. The GM K1 influx rate was ~20% lower in SHRSP rats compared to WKY rats (0.007 ± 0.001 versus 0.009 ± 0.001, p< 0.002, Mann-Whitney) with no significant differences observed in WM (**Fig. 1e**). However, model fit results were more variable in WM when compared to GM. No differences in efflux k2 rates were observed across tissue compartments and strains over the 160 min experimental time window (**Fig. 1f**). Gd-DOTA tracer uptake, as measured by the distribution volume V(t), was 23% and 10% lower in GM and WM, respectively, in SHRSP when compared to WKY rats (**Fig. 1g**). Kinetic 1-compartment modeling applied to data extracted from pre-selected brain regions of interest failed (as evaluated by ‘goodness-of-fit’ parameters) and regional GS transport differences across strains could not be explored using this approach.

### rOMT framework for tracking advection and diffusion modes of GS transport

The rOMT formulation with the inclusion of both advection and diffusion terms in the constraint (continuity) equation is described in detail in the Methods section. Here we highlight that the Lagrangian formulation was used to construct so-called ‘pathlines’ for visualizing GS transport flows in one comprehensive figure, derived from the rOMT returned velocity field and interpolated images (see methods and **Supplementary Fig. 2**). As such, a Lagrangian pathline traces the trajectory of a specific particle over a pre-defined time interval. A *particle* may refer to a parcel of mass or an individual substance. For our purposes, we use *particle* interchangeably with *solute*. Observations of particle attributes, such as ‘speed,’ are made from the vantage point of the particular particle as it traverses its trajectory. In this way, ‘pathlines’ and ‘speed-lines’ encapsulate the moving particle dynamics in the GS network in a single entity. Additional information, such as flow volume (size of the pathline network), reflecting the total flux of particles, is extracted from the pathlines and used in the following for analyzing GS transport across tissue compartments and brain regions. For ease of narration, we refer to the extraction and visualization of the ***La***grangian representation of ***G***lymphatic ***D***ynamics as our ***GLaD*** framework (**Supplementary Fig. 2**). rOMT+GLaD analysis was performed on DCE-MRI images taken over the 120 min interval starting at the time of peak signal (**Supplementary Fig. 2a**). This time interval afforded the best representation of GS transport because the signal-to-noise ratio is maximized (peak) and redistribution of Gd-DOTA tracer into whole brain parenchyma is maximized.

Here we also note that the rOMT algorithm captures the Fick’s law aspect of the flow model, namely that the diffusion mode of particle transport is controlled by the gradient of the concentration. This was accomplished using a custom built ‘diffusion’ phantom incorporating the microdialysis technique^44^ (**Supplementary Fig. 3** and **Movie 1**). The rOMT model was applied to the DCE-MRI data acquired in the diffusion phantom. Knowing that the solute transport in the phantom was solely diffusive, we examined how our model performed if we forced fictitious advective transport. This was done by using very low diffusivity values that were intentionally insufficient to mimic the diffusing intensity patterns observed in the DCE-MRIs. This numerical intervention thereby forced advection to compensate for the lack of diffusion in the modeling of interpolated images. Although the advective and diffusive fluxes influence each other (see Methods), the advective flux is primarily influenced by time-varying changes in voxel intensity while the diffusive flux is primarily influenced by spatial variations. Attempting to account for both temporal and spatial changes in intensity with advection alone resulted in an irregular flow field in the interpolated images. It therefore was not surprising that efforts to extract meaningful (smooth) pathlines with our aforementioned GLaD framework failed on the phantom data under these conditions. Next, we ran the rOMT model on the DCE-MRI diffusion phantom data again, this time with an appropriate diffusivity value (experimentally determined). The resulting flow field was governed by the diffusive flux, as expected. As shown (**Supplementary Movie 1**), diffusive flux vectors point from voxels with higher intensity toward voxels with lower intensity and there is near-perfect consistency between the acquired DCE-MRI images (**Fig. 2a**) and the returned interpolated model images (**Fig. 2b**). This numerical experiment proved informative for understanding the advantages of adding the diffusion term to the classical dynamic OMT formulation and the roles of each transport mode (see methods).

**Figure 2:**
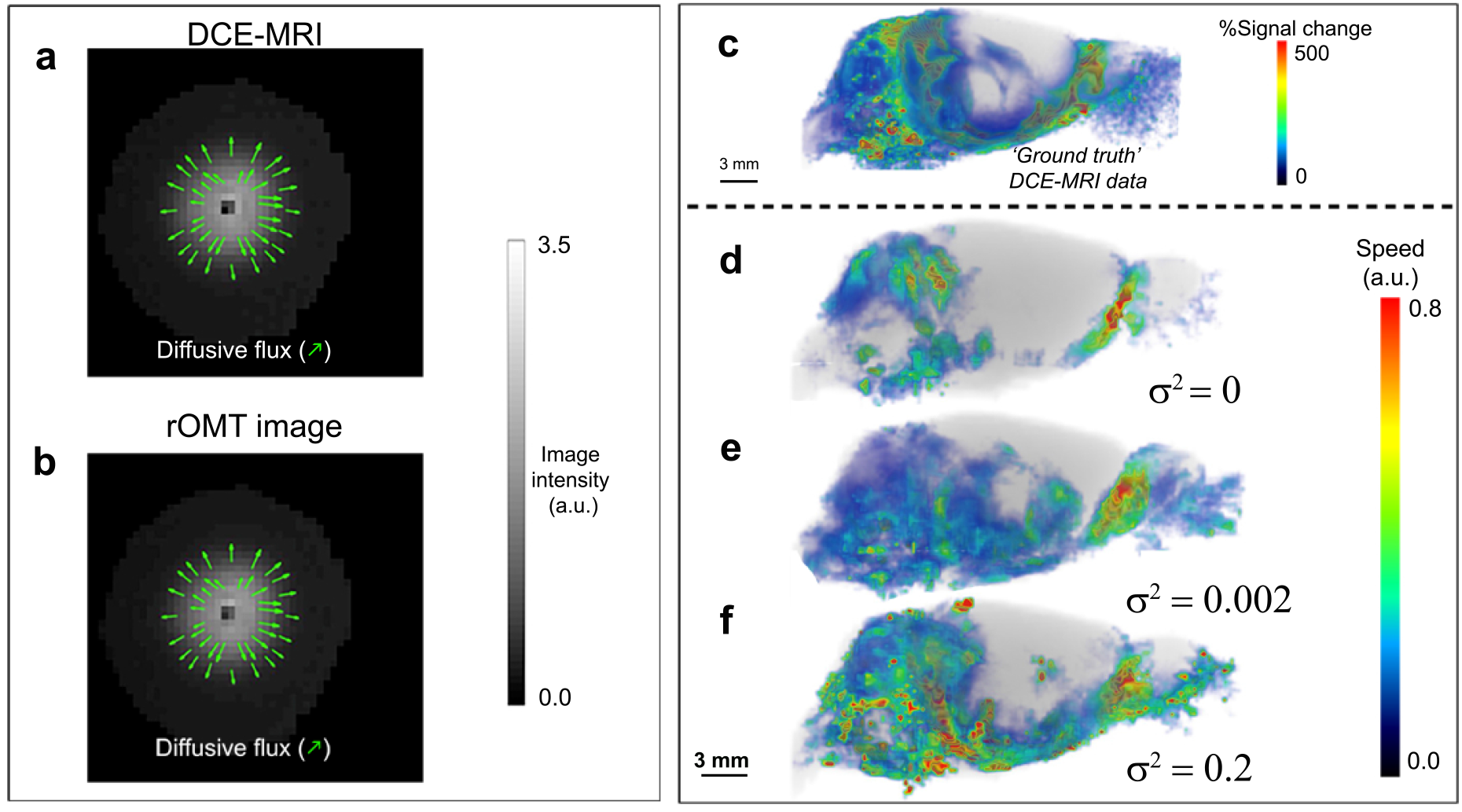
Diffusion is required for rOMT model data to capture in vivo GS transport patterns. **(a)** Agar distribution after ~ 2 hrs of diffusion, as captured with contrast-enhanced MRI, is shown in grayscale. The diffusive direction, proportional to the negative of the image gradient, is shown by the green arrows. (**b)** The corresponding rOMT-returned image is shown in grayscale with diffusive flux vectors characterizing the flow behavior and in alignment with Fick’s law are overlaid in green for comparison to ‘ground truth’ (a). **(c)** Whole brain % signal from baseline at 100 min after contrast injection is shown for context. Corresponding color-coded flow speeds along streamlines computed from the rOMT-derived vector field associated with 100 min after contrast injection for multiple diffusion levels are compared, illustrating how an appropriate diffusion value was selected. **(d)** With no diffusion, the flow shows exaggerated peripheral advective behavior and central parenchymal transport is unaccounted for. **(e)** Using moderate but sufficient diffusion yields feasible peripheral behavior and accounts for transport in deeper parenchymal regions. (**f)** With high diffusion, more activity is accounted for but exaggerated peripheral behavior is also seen.

### Diffusion and advection constraints are both required for matching an rOMT model

Adding diffusion in the rOMT model was required for consistency between GS transport patterns observed in the live rat brain (**Fig. 2c**) and is one of the key features of rOMT transport. In order to choose an appropriate diffusivity value, we tested multiple values and examined how the flow fields changed. Specifically, we computed streamlines from the rOMT derived velocity fields at each time step along with the corresponding speeds. Streamlines are curves that are tangent to the velocity field at a fixed time, informing the collective instantaneous behavior of the flow. We chose this approach over the GLaD-pathline analysis in order to investigate the ‘smoothness’ of the flow field at individual time steps, which should be directly affected by diffusion. A representative example of speed (color-coded) from a WKY rat along the streamlines is shown in **Figs. 2d-f** illustrating the effect of increasing the strength of the diffusion term ‘σ^2^’ in the rOMT algorithm. With either no diffusion (**Fig. 2d**) or too much diffusion (**Fig. 2f**), the captured peripheral behavior is dubiously overactive while information about the flow in the deeper regions, where parenchymal transport of interest occurs, is altogether unrealized. We determined that an intermediate amount of diffusion (σ^2^ =0.002) was required (**Fig. 2e**) in order to replicate observed GS transport in both peripheral and central regions (see the Methods section for a more detailed discussion). This led us to conclude that parenchymal GS transport is governed by both advection and diffusion. We also note that future work will entail improvements in the diffusion model incorporating, for example, spatially varying diffusion as new developments in the detailed understanding of diffusion in tissue becomes available.

### Solute speed varies across tissue compartments

To derive GS solute transport across tissue compartments at the voxel-level, we applied the proposed rOMT+GLaD analytical pipeline to segmented whole brain DCE-MRIs acquired in WKY and SHRSP rats. Examples of pathlines color-coded for speed derived in CSF, WM and GM compartments from a WKY and a SHRSP rat are shown in **Figs. 3b**, **and 3c**. Anatomically, pathlines in GM were more prominent in the cerebellum, hippocampus and hypothalamus of WKYs (**Figs. 3b**). In WM of WKY rats, pathlines dominated in the brainstem and midbrain (**Figs. 3b**). In SHRSP rats, subarachnoid CSF pathlines showed overall more heterogeneously distribution and were most prominent on the ventral surface (**Fig. 3c**, **top**). In WKY rats, total pathline volume in the CSF compartment (representing the total CSF flux of ‘moving’ solutes over the 120 min selected time interval) was 20% higher when compared to SHRSP rats (7124 ± 605 voxels vs 5822 ± 636 voxels, p=0.001, Mann-Whitney, **Fig. 3d**), suggesting that more CSF was available to enter the GS in WKYs and that CSF solute flow was more restricted in SHRSP rats. In SHRSP rats, the lower than normal CSF pathline volume was mirrored by reduced pathline volume in both GM and WM compartments when compared to WKYs (**Fig. 3d**) signifying overall decreased GS transport in the hypertensive rat stain. In the corpus callosum, pathline volume was variable with a trend towards decreased solute transport in the SHRSP compared to WKY rats (582 ± 236 vs 330 ± 120, p=0.051, Mann Whitney).

**Figure 3:**
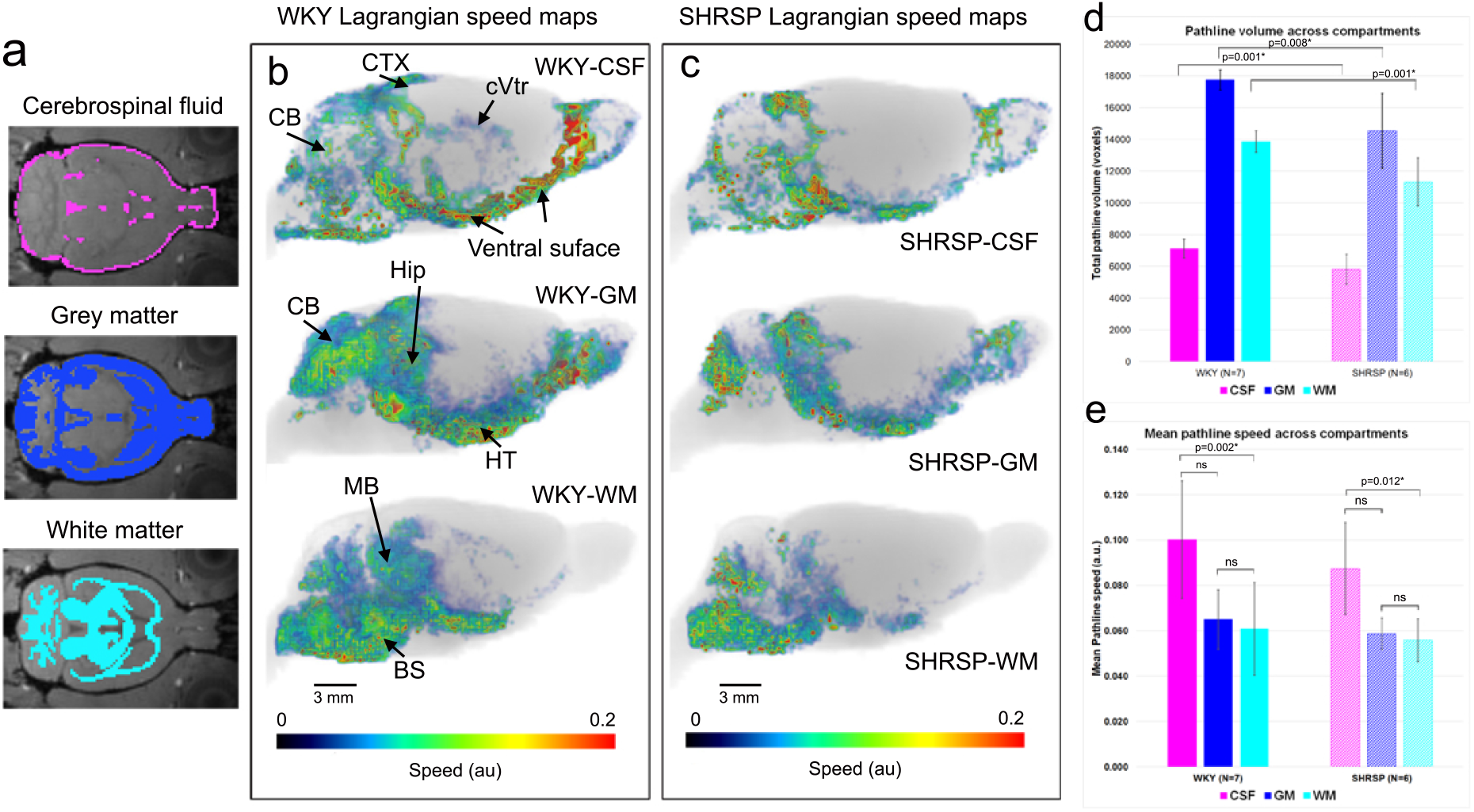
rOMT whole brain analysis reveals varying solute speed across tissue compartments. **(a)** Cerebrospinal fluid (CSF), gray matter (GM) and white matter (WM) masks overlaid on the brain anatomical template to illustrate the spatial distribution of the brain tissue compartments used for the rOMT analysis. **(b)** Pathlines – color-coded to denote speed – overlaid on corresponding anatomical brains are shown from a WKY rat in CSF, WM and GM compartments. Red and blue colored pathlines denote fast and slow pathline speeds, respectively. In the CSF compartment, pathlines are fast moving at the ventral surface. Pathlines with slower speed lines can be seen inside the cerebral ventricles (cVtr). Speed pathlines in GM and WM compartments are variable: faster speed in the periphery and slower speed profiles as they penetrate into deeper parts of parenchyma. CTX=cortex; CB=cerebellum; Hip=hippocampus; HT=hypothalmus; MB=midbrain; BS=brain stem. **(c)** Corresponding pathlines in the three tissue compartments from a SHRSP rat are shown. It is clear that pathlines in CSF are sparser and more heterogeneously distributed in the SHRSP rat when compared to the WKY rat. For example, there are no pathlines above cortex and very few superficial to the cerebellum. For SHRSP, there are also less pathlines in the GM and WM, as compared to the WKY rat. **(d)** Analysis of the total flux of solutes in the pathline network (number of voxels) revealed decreases in all three compartments in SHRSP when compared to WKY rats. **(e)** In both WKY and SHRSP strains, mean pathline speed in CSF was significantly faster than mean pathline speed in the WM compartment. Data are mean ± SD. Mann-Whitney’s test was used for cross-strain analysis. Friedman’s ANOVA was used for within-strain regional analysis. *significant at level alpha=0.05.

We next analyzed the time-varying particle attributes associated with the pathlines, that is, ‘speed’ given by the magnitude of the transport vectors derived by the rOMT+GLaD procedure. Speed trajectories of the pathlines are displayed as color-coded maps for each tissue compartment (**Figs. 3b, c)**. In WKY rats, fast speed (red color) was evident in the CSF subarachnoid compartment on the ventral surface of the brain, superficial to the cerebellum and cerebral cortex (**Fig. 3b**, **top**). Slower (blue color) speed trajectories were observed inside the cerebral ventricles (**Fig. 3b top**). In WKY and SHRSP rats, mean pathline speed was ~40% faster in CSF when compared to the GM and WM compartments, but significant differences were documented for WM only (**Fig. 3e**). Differences in solute speed between CSF and parenchyma can be explained by 1) more restricted transport in the smaller, 15-20% ISF space^45^ when compared to non-cellular CSF spaces given that the solute ‘Gd-DOTA’ is an extracellular tracer in normal brain; 2) less overall advection as the solute moves away from larger pulsatile pial vessels; 3) increased diffusion mode of solute transport in deeper portions of parenchyma. Mean pathline speed in CSF of SHRSP was within a similar range to those of WKY rats (0.10 ± 0.026 versus 0.09 ± 0.02, p=0.558) and no differences in mean speed were observed in GM and WM compartments between the two strains. We also compared *directional* tracer movement from CSF into the tissue compartment between WKY and SHRSP rats (**Supplementary Fig. 4**). Using the rOMT-derived velocity field, we evaluated the total (advective + diffusive) flux across this surface with attention to the net mass moving *from* the CSF compartment *into* the tissue compartment. This analysis showed that the average net mass transported into the tissue compartment was significantly higher for WKY when compared to SHRSP rats (**Supplementary Fig. 4**).

### Solute speed varies across brain regions

We next considered regional GS transport using GLaD analysis applied to the dynamic DCE-MRI data extracted from pre-selected brain regions distributed in the GS network including the hippocampus, cerebellum, superior colliculus and basal forebrain regions (**Fig. 4a**). Using the same visualization scheme as for the whole brain data, **Fig. 4** shows pathlines color-coded for speed captured inside cerebellum, superior colliculus, hippocampus and basal forebrain from a representative WKY (**Figs. 4b**) and SHRSP rat (**Figs. 4c)**. In the SHRSP rats, the total volume of pathlines was significantly reduced in the cerebellum, hippocampus and superior colliculus when compared to WKY rats (**Fig. 4d**). Furthermore, GS transport varied across brain regions as evidenced by the varying distribution of pathlines across the brain in both strains. In the SHRSP rats, pathlines were sparser some areas (e.g., hippocampus) than others (e.g., basal forebrain) possibly due to chronic hypertension (**Fig. 4d**). For both WKY and SHRSP rats, it was also evident that pathline speed was not uniform across regions (**Fig. 4e**). Within normal WKY rats, mean pathline speed in the basal forebrain region was significantly increased when compared to the hippocampus (0.102 ± 0.025 vs 0.055 ± 0.016, p=0.006, Friedman test) and superior colliculus (0.102 ± 0.025 vs 0.053 ± 0.012, p=0.021, Friedman test). Within the SHRSP rats, mean pathline speed also varied across regions (**Fig. 4f**). Across strains, the analysis revealed that mean speed in the superior colliculus was significantly reduced in the SHRSP when compared to WKY rats (0.053 ± 0.012 vs 0.040 ± 0.005, p=0.022, Mann Whitney).

**Figure 4:**
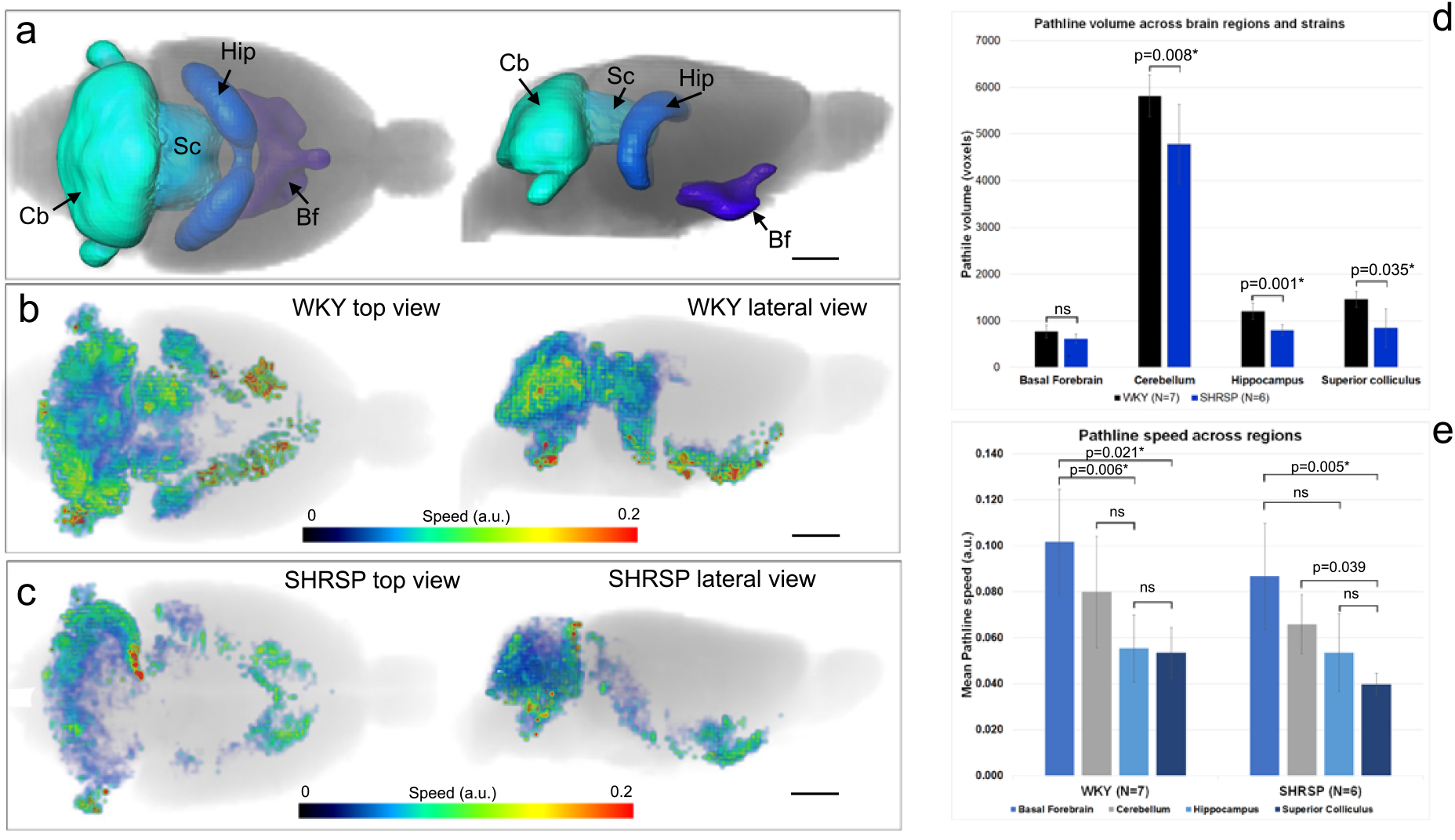
Solute speed is not uniform across brain regions. **(a)** Volume rendered anatomical delineation of brain regions including the cerebellum (cb), superior colliculus (Sc), hippocampus (Hip) and basal forebrain (Bf) that were used for the analysis. Top and lateral views of the regions are shown. **(b)** Color-coded pathlines depicting speed captured over 120 min are shown from a normal WKY rats inside the regions of interest. In the normal WKY rat, speed trajectories in the cerebellum penetrate the entire region. Pathlines at the level of the dorsal hippocampus are sparser when compared to the ventral hippocampus. **(c)** Color-coded pathlines depicting speed captured over 120 min are shown from a SHRSP rat for the same regions. Note that in this particular SHRSP rat, pathlines barely reach the superior colliculus and are not observed in the center of the cerebellum. **(d)** The total pathline volume for each region was calculated and compared between WKY and SHRSP rats. Except for the Bf, all regions showed lower pathline volume in SHRSP when compared to WKY rats. **(e)** Mean pathline speed varied across brain regions in both strains. The fastest mean speed was observed in the basal forebrain for both strains. Data are mean ± SD. Mann-Whitney’s test was used for cross-strain analysis. Friedman’s ANOVA was used for within-strain regional analysis. *significant at level alpha=0.05.

### Solute transfer from PVS-to-tissue is impaired in SHRSP rats

To further explore GS solute transport dynamics between the two strains, we acquired DCE-MRIs at higher spatial resolution using a smaller 1-cm RF-surface coil positioned above the left hemisphere of the rat’s head to capture transport in the PVS of pial arteries at the circle of Willis and along the entire MCA traveling across the left hemisphere (**Movie 2**). To boost signal-to-noise in the PVS, we administered 20 μL of 1:5 Gd-DOTA diluted with sterile water at a rate of 1.5 μL/min into the CSF. The higher Gd-DOTA levels in CSF enabled us to track solute transport along the entire left hemispheric MCA at a voxel resolution of 0.15×0.15×0.15 mm^3^. We first extracted time signal curves (TSC) from the PVS along the MCA on the ventral surface (a.k.a. ‘MCA-PVS root’, **Fig. 5a**) and observed strikingly different signal dynamics between SHRSP and WKY rats (**Fig. 5b**). While the MCA-PVS root TSCs from SHRSPs and WKYs were characterized by similar time-to-peak and peak magnitude, the signal ‘relaxation’ phase was faster in WKYs, suggesting faster transit of solutes into parenchyma compared to SHRSP rats (**Fig. 5b**). This interpretation was supported when comparing DCE-MRIs at the base of the brain after ~3 h of CSF Gd-DOTA circulation (**Figs. 5c, d**). Significantly higher Gd-DOTA signal was evident along the root PVS-MCA of SHRSP rats (**Fig. 5d**) but not along the MCA distal to this point when compared to WKYs (**Figs. 5e**) suggestive of physical obstruction (or ‘slow-down’) of MCA-PVS flow in this area.

**Figure 5:**
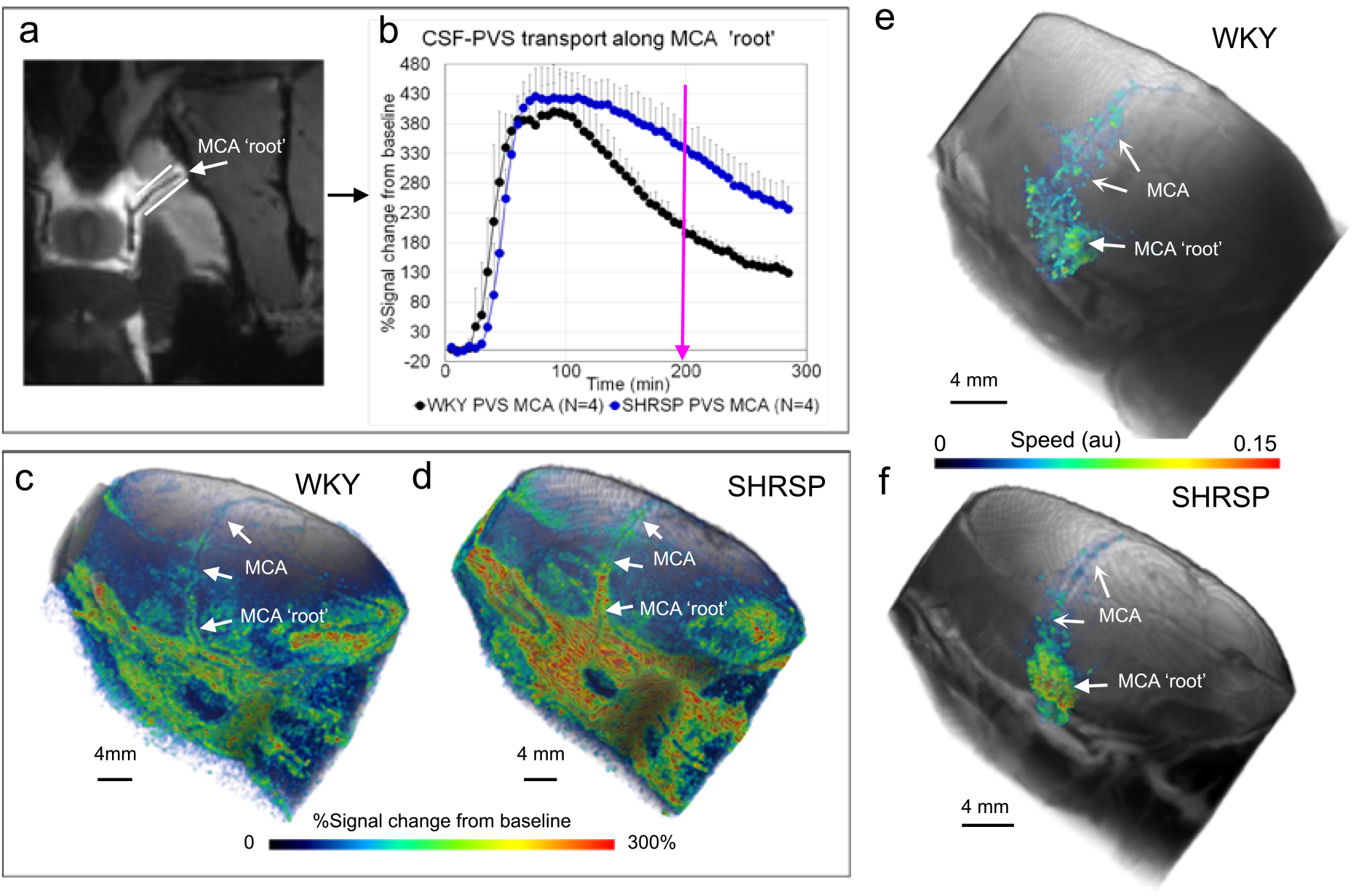
Solute transfer from the perivascular space (PVS) along the middle cerebral artery. **(a)** Contrast enhanced MRIs from a WKY rat at the level of the circle of Willis near the take off of the middle cerebral artery (MCA). Contrast in the perivascular space along the circle of Willis and MCA is evidenced as a high intensity signal along the vasculature, which is black. Regions of interest (ROI) along the MCA (white lines) were used to capture time signal curves (TSC) from the dynamic contrast enhanced (DCE) MRIs acquired in WKY and SHRSP rats. **(b)** Mean TSC extracted from WKY (N=4) and SHRSP (N=4) rats along the first segment of the MCA (‘root’) are displayed. Data are mean ± SD. It is evident that while ‘time-to-peak’ and peak magnitude are within same range for the two strains, the signal decline (representing redistribution of contrast from PVS to tissue and tissue clearance) is strikingly faster in WKY when compared to SHRSP rats. **(c)** Volume-rendered, color-coded signal map after ~180 min of circulation of CSF contrast in a WKY rat overlaid onto the corresponding anatomical template. High and low contrast signal amplitude is given by red and blue colors respectively. Contrast is observed along the entire MCA during its trajectory across the left hemisphere. **(d)** Corresponding color-coded signal map from a SHRSP rat showing much higher levels (red color) of contrast in the PVS along the circle of Willis and along the MCA root. Heterogeneously distributed contrast along the MCA is more evident in the SHRSP when compared to WKY rat. **(e-f)** The color-coded speed maps along the MCA are overlaid on the anatomical volume rendered MRIs of the left hemisphere of a WKY **(e)** and SHRSP **(f)**. While pathline speed appeared to be homogeneous along the MCA in WKY rats, pathline speed in SHRSP rats was high in areas localized to the ‘root’ area and lower distal to this point.

The rOMT+GLaD pipeline was applied to extract pathline speed information along the MCA. Pathline start points were selected from binary masks capturing tissue along the MCA in order to analyze transport behavior stemming from this specific region (see Methods). The color-coded speed maps along the MCA are overlaid on the anatomical volume rendered MRIs of the left hemisphere (**Figs. 5e, f**). While pathline speed appeared to be homogeneous along the MCA in WKY rats (**Fig. 5e**), pathline speed varied along the MCA in SHRSP rats (**Fig. 5f**). Specifically, in SHRSP rats, pathline speed was increased by 3-fold along the MCA ‘root’ when compared to speed along the MCA distal to this area (0.12 ± 0.04 vs 0.04 ± 0.01, p=0.029, Student’s t test). Corresponding measurements in WKY rats did not reveal a significant differences in speed along the MCA (p=0.25). We also noted that in WKY pathlines had migrated >4 mm away from the MCA-PVS into the adjacent tissue (**Fig. 5e**). In contrast, in SHRSP rats, pathlines were observed in closer proximity to the PVS-MCA (**Fig. 5f**).

We also performed directional analysis of the advective flux direction along the MCA in normal WKY rats (**Fig. 6**). Advective flux vectors were calculated across the boundaries of the binary mask created along the MCA from the rOMT output. **Fig. 6** shows advective flux vectors (normalized to have unit length) along the MCA overlaid on the anatomical MRI from a WKY (**Figs. 6a, b**) and SHRSP (**Figs. 6c. d**). **Figs. 6a-d** highlight directions of the advective flux vectors from the ventral surface of the brain at the level of the circle of Willis from where the MCA take off (white boxes in **Figs. 6a, c**). In the WKY rat the advective flux vectors’ direction is following the path along the MCA in a continuous pattern (**Figs. 6 a, b**). In the SHRSP rat, the advective flux vectors are following the direction of the MCA part of the way, but then the flow pattern become disrupted and advective flux vectors can be seen diverging from the MCA flow direction (**Figs. 6c, d**). The deflections in advective flux vector directions along the MCA in SHRSP compared to WKY rats is also clearly appreciated in ‘caudal’ projection views (**Figs. 6e, f**).

**Figure 6:**
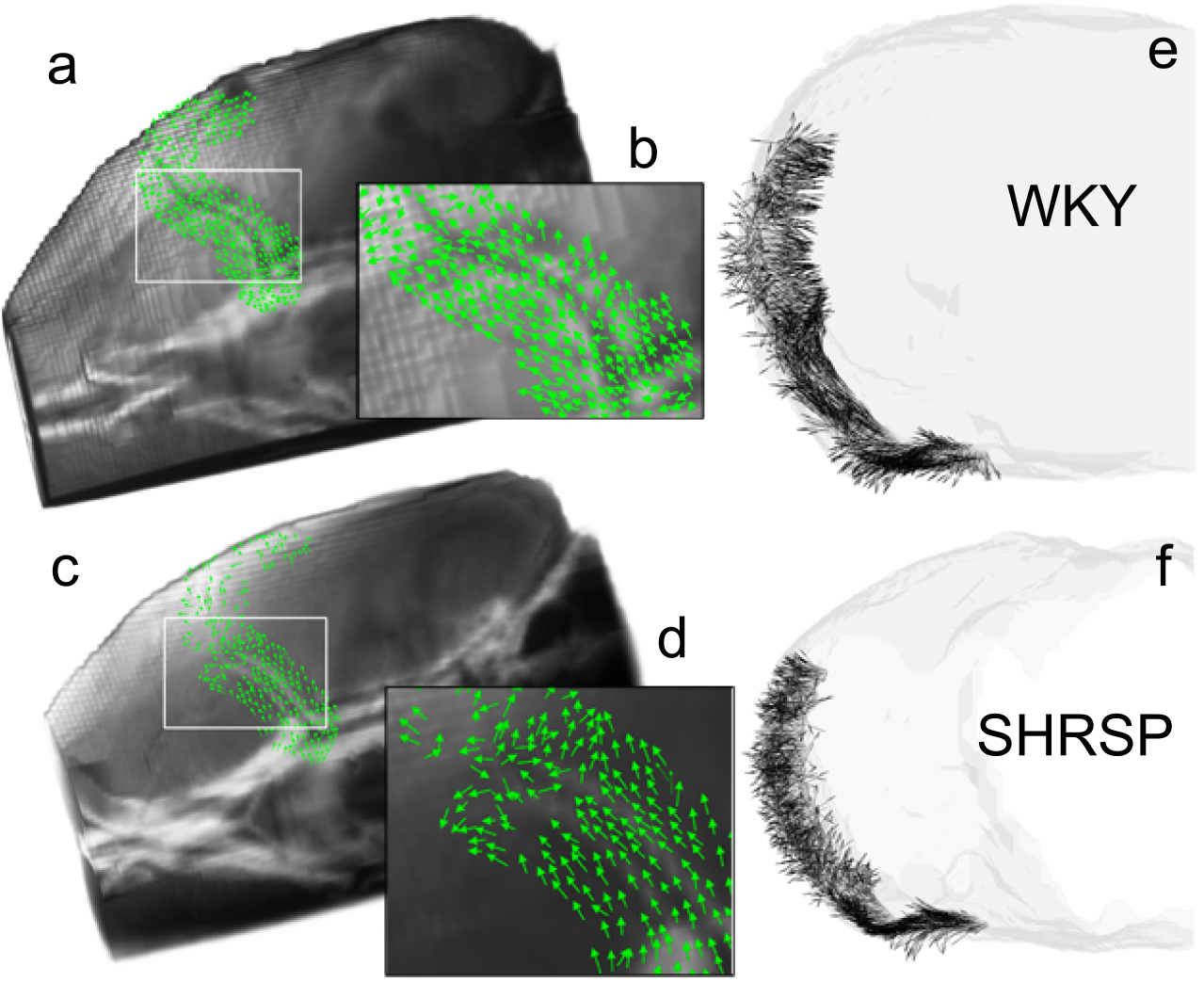
Advective flux vectors along the MCA show aberrant flow in SHRSP. **(a, b)** Advective flux vectors across the boundary along the MCA, determined from the binary mask, were calculated from the rOMT output. The advective flux direction, determined by normalizing the advective flux vectors to have unit length, are shown overlaid on an anatomical mask of a WKY rat. The ventral surface at the level of the circle of Willis and MCA origin is highlighted. In the WKY rat, the directions of the advective flux vectors follow the path along the MCA in a continuous pattern. **(c, d)** In a corresponding view of a SHRSP rat, the advective flux vectors are pointing in the same direction along the first segment of the MCA along the ventral surface. However, distal to this point, the vectors start diverging away from the MCA. **(e, f)** The deflections in the direction of the advective flux vectors at the transition from ventral to lateral segments of the MCA in the SHRSP rat compared to the WKY rat is also clearly visible in ‘caudal’ projection views.

### Mechanisms underlying the decreased GS transport chronic hypertension

We next explored major drivers of GS transport to uncover possible mechanisms underlying the differences observed between SHRSP and WKY rats. Increases^19^ and decreases^1, 20^ in vascular pulsatility have been shown to accelerate and decelerate GS influx, respectively. Only subtle differential effects of rat strain on the mean heart rate recorded continuously during the glymphatics MRI experiments were noted (**Supplementary Table 1**). In fact, mean heart rate was slightly increased in SHRSP compared to WKY rats, leading us to conclude that changes in heart rate were unlikely to contribute to GS transport impairment observed in the SHRSP rats. In a series of randomly selected awake, non-anesthetized rats from age-matched cohorts of WKY and SHRSP rats, non-invasive blood pressure was measured using the tail-cuff method. Systolic and pulse pressures were documented to be significantly higher in SHRSP compared to WKY rats (**Supplementary Table 1**). General anesthesia will dampen blood pressure when compared to the? awake state and SHRSP used for the glymphatics experiments may not have been characterized by higher blood pressure when compared to WKY rats. Therefore, in a separate series of rats with femoral artery catheterization, we documented that mean arterial blood pressure indeed remained higher in SHRSP even under anesthesia when compared to WKY rats (**Supplementary Table 1**). Based on these data, we concluded that higher than normal pulse pressures in SHRSP rats might have contributed to the impaired GS transport observed in our study.

State of arousal^46^ and type of anesthesia^47^ have also been shown to affect GS transport. We did not measure electroencephalogram in the experiments, however anesthetic depth was evaluated indirectly by quantifying anesthetic drugs administered and was found to be similar between strains (**Supplementary Table 1**). We therefore concluded that differences in depth of anesthesia are unlikely to have contributed to the GS transport alterations observed.

Polarized perivascular expression of AQP4 water channels has been associated with rapid transport of small molecular weight solutes from PVS to brain ISF^1, 36, 48^. We preformed immunohistochemistry on a smaller series of WKY (N=4) and SHRSP, (N=4) to examine the AQP4 expression patterns associated with small vessels and capillaries in select brain regions. In WKY rats, the distribution of AQP4 immunofluorescence appeared homogeneously distributed in the occipital cortex (**Fig. 7a**) and ventral hippocampus (**Fig. 7c**). In SHRSP rats, increased immunofluorescence was noted in patchy areas associated with penetrating cortical and hippocampal vessels (**Figs. 7b, d**). AQP4 immunofluorescence intensity was measured along linear ROIs crossing the lumen of small vessels or cerebral capillaries and converted into a polarization index (**Supplementary Fig. 5** **and methods**). This analysis showed that AQP4 immunofluorescence surrounding the small vessels and capillaries was more polarized in WKY (higher polarization index) when compared to SHRSP rats (**Figs. 7g, h**). We also executed qPCR to measure the mRNA expression levels of GFAP and AQP4 in a separate series of WKY (N=8) and SHRSP (N=8) rats in the following regions: cerebellum, cortex, hippocampus, striatum and thalamus. This analysis revealed no statistical differences in AQP4 or GFAP mRNA expression between WKY and SHRSP rats (**Supplementary Fig. 6**). Collectively, based on these data, we conclude that impaired perivascular AQP4 polarization in SHRSP rats (but not increased total tissue AQP4 expression or astrogliosis) might have contributed to the reduction in PVS-to-brain solute transfer observed in the SHRSP rats when compared to WKY rats.

**Figure 7:**
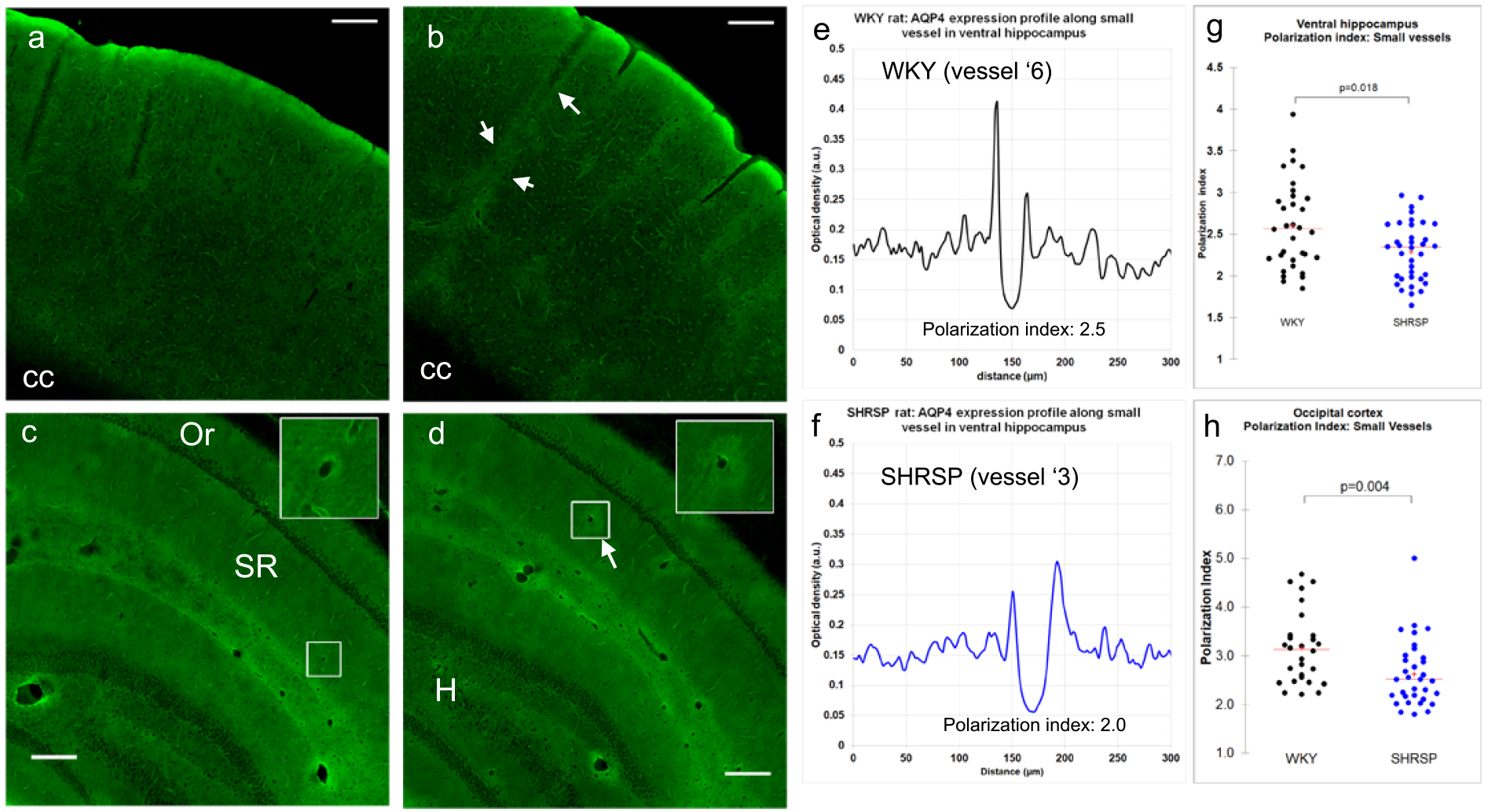
Perivascular AQP4 water channel expression is altered in SHRSP rats. **(a**) Confocal representative micrographs show AQP4 immunoreactivity extending from the occipital cortical surface towards the corpus callosum (cc) and shows that AQP4 was distributed uniformly. **(b)** Equivalent confocal montage from the ventral hippocampus of a WKY rat. AQP4 is distributed homogeneously in the Stratum Oriens (Or) and Stratum Radiatum (SR). **(c)** AQP4 distribution in occipital cortex from a SHRSP rat showing higher intensity expression along penetrating vessels (white arrows). **(d)** In SHRSP rats, intensely expressed AQP4 can be noted in patchy areas surrounding vessels in the SR (arrows) and occasionally in the hilus (H) region. **(e-f)** AQP4 intensity was measured along linear ROIs crossing the lumen of small vessels or cerebral capillaries in WKY and SHRSP rats and converted into a polarization index (**Supplementary Fig. 5 and Methods** for more information). **(g)** Perivascular AQP4 polarization index of small vessels in the ventral hippocampus is significantly reduced in SHRSP when compared to WKY rats. **(h)** In the occipital cortex, the perivascular AQP4 polarization index of small vessels was also significantly reduced in SHRSP when compared to WKY rats. Scale bar=250 μm.

Finally, we explored vascular and structural anatomy at the level of the circle of Willis and area of the MCA root where the contrast in the PVS appeared to be ‘obstructed’. **Supplementary Fig. 7** show representative brain sections from WKY rat and SHRSP rat were immunolabeled with an antibody to collagen IV to identify cerebral vessels and micro-vessels (**Supplementary Fig. 7c**, **insert**). In WKY rats smaller networks of micro-vessels were observed in distinct areas at the base of the brain surrounding penetrating arteries (**Supplementary Figs. 7d-f**). However, in SHRSP rats, corresponding microvascular networks were larger in size and the microvasculature denser (**Supplementary Fig 7g-i**). We speculate that the microvascular network remodeling might have affected the PVS transport from the circle of Willis. More investigations are needed to fully understand mechanisms underlying the ‘obstructed’ PVS flow in SHRSP rats compared to WKY rats.

## Discussion

Our rOMT Lagrangian workflow method applied to DCE-MRI images used to measure GS transport in whole rat brain provides the first evidence of both diffusion and advective solute transport modes in the brain proper. First, in normal brains, rOMT analysis revealed that both advection and diffusion terms were required for pathlines and trajectory speed in the modeled images to resemble GS transport patterns in live brain. Second, particle speed varied strikingly across tissue compartments with faster solute passage in CSF when compared to GM and WM compartments. Third, large differences in regional solute speed within the brain parenchyma were also documented. Specifically, in regions close to the ventral surface near large pulsatile arteries (e.g. basal forebrain) solute speed was 2-fold faster when compared to midbrain and hippocampual regions. Collectively, these novel results demonstrate that advective transport is dominating in the non-cellular CSF spaces and that slower diffusion transport contributes to GS transport as the solute is transferred from the PVS into tissue.

The OMT problem was first posed by Gaspard Monge in 1781^49^ and may be formulated as one of transporting one distribution of mass to another in a manner that minimizes a given cost functional. *Mass* is used in a general sense and originally referred to soil, as Monge was interested in the civil engineering problem of leveling the ground. In our work on the GS, *mass* is represented by signal intensity induced by a paramagnetic contrast solute (previously shown to reflect Gd-DOTA concentrations^50^) administered into CSF and captured by DCE-MRI. The model is related to the Navier-Stokes model but instead of considering momentum, one considers the flow of density^51, 52^. The advection/diffusion rOMT model is simpler to analyze, but nevertheless provides physically meaningful information. Interestingly, there are strong connections between rOMT and the porous media and kinetic models (see the Methods section) previously applied to characterize GS transport^37, 53, 54^. However, in contrast to these models, the rOMT problem does not require *a priori* identification or delineation of boundaries, time-varying input or tissue function, or specification of kinematic parameter values. Other than the assumption of the underlying advection/diffusion equation, no prior assumptions are imposed on the GS flow. Finally, rOMT may be considered to be an optical flow technique^55^, but instead of imposing smoothness on the solutions of the continuity equation, one employs it as a constraint in minimizing a quadratic energy function.

GS dysfunction is reported in neurodegeneration particularly where vascular dysfunction may be involved^7, 35, 36^. We therefore applied the rOMT framework to dissect modes of solute transport changes in an animal model of cerebral small vessel disease (cSVD). cSVD is the major pathology in vascular dementia, and GS dysfunction has recently been highlighted as a novel pathogenetic factor in cSVD^5^. The hypothesized link between GS dysfunction and cSVD is based on the observation of abnormally enlarged PVS in clinical cases of cSVD^56^. We utilized rOMT analysis to characterize modes of GS solute transport across tissue compartments in normotensive WKY and SHRSP rodents. The SHRSP rat is a naturally bred animal with a parent control strain, which mimics human hypertension and cSVD^57^. Another reason that we used the SHRSP rat is that they do not present with hydrocephalus – a drawback inherent to other rat strains with chronic hypertension (e.g. SHR rat^41^) which may affect CSF flow dynamics^7^. The rOMT analysis of DCE-MRI data from the brain showed that the total flux of moving solutes in all tissue compartments (including CSF) was reduced in SHRSP when compared to normotensive WKY rats.

Interestingly, mean solute speed in the CSF and tissue compartments was not significantly reduced in SHRSP when compared to WKY rats. However, a separate analysis of CSF transport in the PVS along the MCA obtained with DCE-MRI acquired with increased spatial resolution and higher contrast-to-noise (increased Gd-DOTA concentration in CSF) revealed altered CSF flow patterns in SHRSP when compared to WKY rats. In WKY rats, solute speed along the MCA was homogeneous whereas in SHRSP rats solute flow became overtly heterogeneous along the MCA. The rOMT analysis of the SHRSP MCA data revealed that solute speed was 3-fold higher at the MCA-PVS ‘root’ area when compared to speed distal to this area suggesting a physical or physiological obstruction at this site.

We investigated possible mechanisms underlying the reduced CSF solute flux in specific regions such as the hippocampus and documented impaired perivascular AQP4 expression in SHRSP compared to WKY rats in hippocampus and occipital cortex which might have contributed to the reduced GS transport patterns. Reduced perivascular expression of AQP4 water channels has been documented in postmortem studies of human cases of Alzheimer’s disease^6^ and in aging rodents^36^.

In summary, the new rOMT Lagrangian framework analysis introduced here established the presence of diffusion and advective solute transport in brain parenchyma supporting the glymphatic system hypothesis and its relevance for waste clearance. In addition, altered solute speed profiles across tissue beds consistent with slower transfer of solutes from PVS to ISF was revealed in chronic hypertension. Future studies should be conducted to further optimize the rOMT Lagrangian analysis developed here, and to confirm the observed reductions in solute transport in human chronic hypertension and cSVD. In addition, future rOMT work will consist of including a source term in our set-up to include physical parameters (e.g., pressure) as well as combining OMT with other metrics such as Fisher-Rao in the cost functional to alleviate the explicit constraint of mass preservation in the model^58^.

## Methods

### Animals

The local institutional animal care and use committees at Yale University, New Haven, USA approved all animal work. 8-month old WKY female rats and aged-matched female SHRSP rats (Charles River, Wilmington, MA, USA) were included in the study. All the animals were given standard rat chow and water *ad libitum*.

### Blood pressure measurements in SHRSP and WKY rats

Non-invasive blood pressure measurements were performed in a selection of awake WKY and SHRSP rats using tail cuff method (CODA monitor, Kent Scientific Corporation, USA). Three measurements were taken × 3 with 5-min interval from each rat and average of systolic and diastolic pressures were used for the analysis. In a separate series of anesthetized WKY (N=6) and SHRSP rats (N=5) mean arterial blood pressure was measured using a catheter inserted into the femoral artery. Anesthesia for these rats were performed with isoflurane and dexmedetomidine as described below.

### Anesthesia and cisterna magna catheter

For all glymphatic experiments, rats were anesthetized with 3% isoflurane delivered into an induction chamber in 100% O2, received dexmedetomidine (0.01mg/kg i.p.) and glycopyrrolate (0.2 mg/kg i.p.). Surgical anesthesia was maintained with 2-2.5% isoflurane and the rats were breathing spontaneously throughout the experiments. A small 5-mm copper tube (0.32 mm o.d., Nippon Tockushukan, MFG. CO., LTD, Tokyo, Japan) attached to a PE-10 microcatheter was implanted into the CSF via the cisterna magna and secured in place using cyanoacrylate glue. Following surgery, the rats were transferred to the MRI instrument. Anesthesia was maintained throughout the experiment by 0.5-1.0% isoflurane in 1:1 Air:O_2_ mixture delivered via a nose cone, and a subcutaneous infusion of dexmedetomidine (0.015-0.020 mg/kg/hr)^59^. During imaging, respiratory rate, oxygen saturation, body temperature and heart rate were continuously monitored using an MRI compatible monitoring system (SA Instruments, Stony Brook, NY, USA). Body temperature was kept within a range of 36.5°C~37.5°C using a heated waterbed.

### MRI imaging

All MRI acquisitions were performed on a Bruker 9.4T/16 magnet (Bruker BioSpin, Billerica MA) interfaced to an Avance III console controlled by Paravision 6.0 software. All rats were imaged in the supine position. For rats used for the whole-brain glymphatic experiments, a 3-cm planar receive surface radio-frequency (RF) coil (Bruker) was positioned under the head of the rat with a separate preamplifier used as a receiver. For rats where glymphatic transport was explored along the middle cerebral artery, a 1-cm planar RF coil (Bruker) was placed above the left side of the rat’s head. A custom-made volume diameter volume coil (50-mm i.d.) was used as a transmit RF coil.

### Glymphatic transport DCE-MRI

Following an anatomical localizer scan, acquisition of the DCE-MRI started using VFA-SPGR sequence. For whole brain glymphatics studies: TR=25 msec, TE=4 msec, 0.30×0.30×0.30 mm FA=15°, number of averages=2, Time/scan=5min). For the MCA glymphatics studies (left hemisphere): TR=25 msec, TE=4 msec, 0.15×0.15×0.15 mm FA=20°, number of averages=1, scan time=4min 10s). A reference phantom filled with 0.1 mM Gd-DOTA was placed in the field of view to allow intensity normalization between each scan as well as allowing receiver gain adjustment. For measurement of GS transport, DCE-MRI were acquired before (3 baseline scans), during and after the infusion of gadoteric acid (Gd-DOTA, DOTAREM, Guerbert LLC, Carol Stream IL) into the CSF via the CM catheter ^53^. Specifically, for whole brain studies we infused 20 μL, of 1:37 Gd-DOTA diluted with sterile water at a rate of 1.5 μL/min. For the higher resolution MCA scans, we infused 20 μL, of 1:5 Gd-DOTA diluted with sterile water at a rate of 1.5 μL/min.

### Voxel-wise morphometric analyses

To study anatomical volumetric differences between WKY and SHRSP rats, we implemented deformation-based morphometry as reported previously^41^. For this purpose, WKY (N=7, 8-months old) and SHRSP (N=6, 8-months old) were scanned using 3D spoiled gradient echo (SPGR) sequence with the following parameters: (TR/TE/FA/Ave=50ms/4ms/7°/4 scanning time= 23min field of view (30×30×15 mm) reconstructed at isotropic spatial resolution of 0.23×0.23×0.23 mm. Morphometric image processing steps were similar to the procedure described previously comprising B1 RF inhomogeneity correction, tissue segmentation, and spatial registration^4^. Following the DARTEL registration algorithm, the voxel-wise Jacobian determinant was calculated for each voxel and subsequently smoothed by a 0.4mm Gaussian smoothing kernel. Voxel-wise statistical analysis was performed by two sample t-tests in the framework of general linear modeling with the total intracranial volume (TIV) as a covariate and statistical significance (p-value) reported in color-coded maps.

### Tissue compartment and ROI specific glymphatic flow analyses

To study tissue compartment specific glymphatic transport, WKY (N=7, 8-months old) and SHRSP (N=7, 8-months old) were scanned using the 3D VFA-SPGR sequence (TR/TE/FA/Ave=15ms/4ms/5~30°/2 scanning time= 30 min 0.30×0.30×0.30 mm) as described previously^41,4^. Proton density weighted (PDW) images calculated from VFA-SPGR were segmented using a custom made SHRSP-WKY rat brain template in deriving spatial registration parameters ^4^. Spatially normalized segmented images were population averaged and thresholded (0.5) to create a population averaged binary mask for GM, WM and CSF, and inverse warped to match the individual rat brain. To study ROI specific glymphatic transport, the publicly available Waxholm Wistar rat brain atlas^60^ was downloaded, and ROIs in the atlas were warped onto the individual SHRSP-WKY rat brains in two steps. First the high resolution T2 weighted Waxholm Wistar rat brain atlas was segmented using the SHRSP-WKY template to derive the registration parameters between the atlas and the SHRSP-WKY template. The registration parameters were then applied onto the ROIs to construct the ROIs in the template space. ROIs in the template space were then inverse warped, which was derived from the tissue compartment analyses, to create the ROIs in the individual brain. Nineteen regions were included in the atlas: anterior commissure, basal forebrain region, brainstem, hippocampus, corpus callosum, descending corticofugal pathway, fimbria of hippocampus, hypothalamus, superior colliculus, cerebellum, cortex, olfactory bulb, penial, periaqueductal grey, septal region, striatum, subiculum, substantia nigra and thalamus. Only four of these (basal forebrain, cerebellum hippocampus and superior colliculus) were used for GS analysis purposes.

### Kinetic analysis of DCE-MRI data from whole brain

Image processing of the series of DCE-MRI data of each rat consisted of head motion correction, intensity normalization, smoothing, and voxel-by-voxel conversion to percentage of baseline signal using SPM12^61, 62^ (https://www.fil.ion.ucl.ac.uk/spm/). The processed images were converted into NIfTI format for further analysis. Brain masks of CSF, GM and WM derived from the morphometric analysis were used to extract compartmental “time-signal curves” (TSCs) of Gd-DTPA-induced contrast changes using PMOD software (PMOD, version 3.8). A 1-compartmental kinetic analysis was used to model the glymphatic movement of Gd-DTPA contrast through brain tissue using the TSCs from the CSF (main input), GM and WM from each rat. Two rate constants (*k*_1_ and *k*_2_) and distribution volume V(t) were calculated from the 1-compartment model using PMOD software (Version 3.8). In the next few sections, we provide the necessary details pertaining to our regularized OMT framework before describing how it was utilized for analyzing the DCE-MRI data.

### Regularized OMT (Eulerian framework)

The ***regularized OMT*** (rOMT) problem^63^ considers the minimization of the energy functional

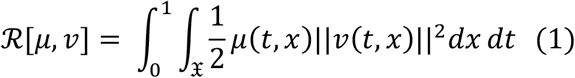

overall time-varying densities *μ* = *μ*(*t*, *x*) ≥ 0 and sufficiently smooth velocity fields *v* = *v*(*t*, *x*) ∈ ℝ^*n*^ subject to the advection/diffusion (constraint) equation

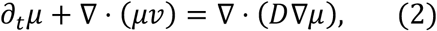

for all *x* ∈ 𝖃, a connected Euclidean subdomain of ℝ^*n*^, over the normalized time interval *t* ∈ [0,1] with initial and final conditions

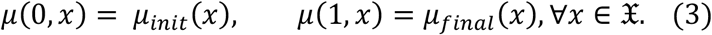

Here, *μ*_*init*_ and *μ*_*final*_ are given observed states of the density at respective times *t* = 0 and *t* = 1, and *D* denotes a symmetric positive definite diffusivity matrix, which may be spatially dependent or independent. When *D* = 0, the constraint (2) is simply the advection equation and we recover the classical dynamic-OMT formulation^34^

From the Calculus of Variations^34, 63^, the Euler-Lagrange equations associated with the regularized OMT problem (1)-(3) shows that the optimal velocity, *v*_*opt*_, has the form

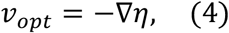

where *η* is the Lagrange multiplier of constraints (2), (3) and satisfies the diffusion Hamilton-Jacobi equation

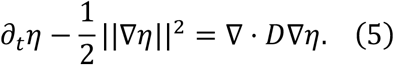

The diffusion term plays a dual role. First of all, it regularizes the flow. In fact, it makes the problem equivalent to the so-called Schrödinger bridge problem, and therefore to a certain type of regularization (related to Fisher information) of the cost function^34, 63^. This leads to smoother flow fields. Secondly, it captures a key feature in modelling the underlying dynamics that may best be understood via compartmental models that will be described in the next section.

### Regularized OMT and compartmental models

Compartmental models^64^ are comparable to spatially averaging the regularized OMT continuity equation (2). More precisely, one may average the continuity equation by integrating it over a control volume (i.e., compartment) *C*:

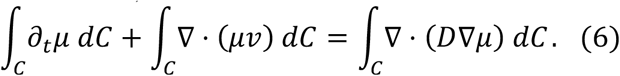

Considering the advective term, we see that

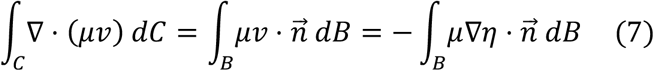

by (4) and the divergence theorem, where 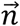 is the unit outward normal of *B*, the compartment boundary. In this form, the OMT-advection may be interpreted as an expression of Darcy’s law, which relates flow velocity to the pressure gradient, as used in the porous media model. We note that the OMT-velocity does not explicitly account for pressure differences while the porous-media-velocity does. One can supplement a pressure dependent source term in the continuity equation, an extension we plan to explore in future work. Considering the unresolved true physical parameter values, this exemplifies the OMT framework’s utility for capturing, representing and informing on meaningful dynamic behavior implicitly encoded in the given observed density distributions, along with interesting insight into how the method works.

Applying the divergence theorem to the diffusive term we find:

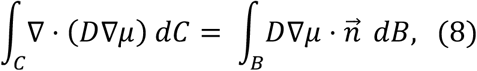

and thus, we see that diffusion is influenced by the density gradient, in accordance with Fick’s law. Noting that the total mass in *C* at time *t* is given by

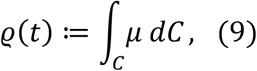

the averaged form of the diffusion-OMT constraint over the given compartment *C* relates the change in its density over time to the flux across its boundary, and (6) may be written as

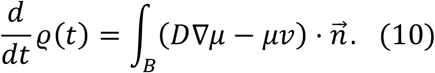

Here, *j*_*AD*_ = −(*D*∇*μ* − *μv*) is the total flux, comprised of both advective and diffusive components. We note that the two are not independent of each other, as is seen by (5).

To illustrate the compartmental regularized OMT formulation, we compare tracer movement from CSF into the tissue compartment between normotensive (WKY) and hypertensive (SHRSP) cases. As such, we take the tissue compartment to be our control volume *C* and consider the surface boundary *B* between the CSF and tissue compartment (**Supplementary Fig. 4a**). For every boundary point *x* ∈ *B*, we can define the unit normal vector 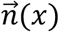 that is orthogonal to the surface at *x* and we orient the surface so that 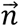 points from the CSF compartment into the tissue compartment (**Supplementary Fig. 4b**). Using the OMT-derived velocity field and smooth images, we can then look at the total (advective + diffusive) flux *j*_*AD*_ across this surface at each time step by considering the component of the flux in the normal direction. At a given time, (10) describes the net mass flowing through the surface. However, we are specifically interested in the net mass that moves *from* the CSF compartment *into* the tissue compartment. We are therefore interested in the normal component of the flux such that 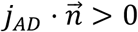. Due to the directional information provided by the rOMT, this can be readily be determined. Fluctuations in voxel-level information given by rOMT are clearly lost when averaged, indicative of the well-known challenge for the kinetic analysis approach for local insight.

### Regularized OMT-Lagrangian framework

The Eulerian framework records measurements at fixed locations for multiple points in time, informing on the collective instantaneous behavior of the flow. We now describe our Lagrangian framework for expressing the trajectories of fixed parcels or particles. In this work we have assumed that the diffusion is constant, *D* = *σ*^2^*I*. While this assumption is somewhat restrictive, it nevertheless gives physically meaningful results. All the theory goes through in the non-constant spatially dependent setting. In future work, we plan to examine this extension in more detail. We note that

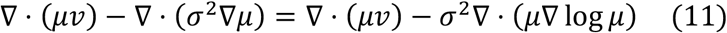

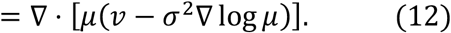

Defining the augmented velocity

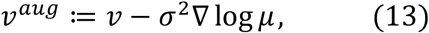

we derive the conservation form of the regularized constraint (2),

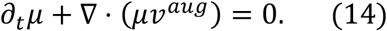

The Lagrangian coordinates *L* = *L*(*t*, *x*) of the optimal trajectory for the regularized OMT system are given by

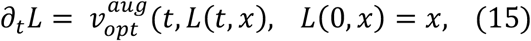

where

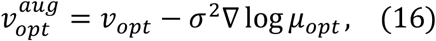

and *v*_*opt*_ and *μ*_*opt*_ denote the optimal arguments of the regularized OMT problem (1)-(3).

### Numerical regularized OMT model for glymphatic system

In this section, we discuss the considerations taken for applying the regularized OMT model to the glymphatic system and its numerical implementation. For our purposes, MR signal enhancement induced by Gd-DOTA is treated as density (the relationship between mass and signal enhancement has been previously validated^39^). The images then provide discrete observations *μ*_*init*_ and *μ*_*final*_ of the tracer distribution at some initial time *t* = 0 and a final time *t* = 1, respectively. Space is discretized as a cell-centered grid of size *m*_1_ × *m*_1_ × *m*_3_ with a total number of *M* cells, each with uniform length *dx*, as delineated by the voxels. Time is split into *N* intervals of length *dt* and the *N* + 1 time steps are denoted by the subscript *n* where *n* = 0 corresponds to the time *t* = 0, *n* = *N* corresponds to the time *t* = 1 and *dt* * *N* = 1. Bold font is used to denote linearized variables and we use 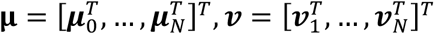 to represent the temporal concatenation of the linearized densities and velocities, respectively. Note that *v*_*n*_ is the velocity field that describes the evolution between *μ*_*n*−1_ and *μ*_*n*_ subject to the advection/diffusion equation (2), for *n* = 1, …, *N*. Following the work of^65^, the numerical method used here was chosen for its nice properties for differentiating and optimizing the resulting discrete regularized OMT problem, which will become clear in the discussion to follow.

We start by discretizing the advection/diffusion equation (2) as

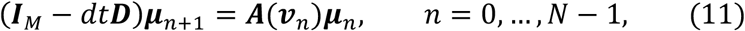

where ***I***_*s*_ is the *s* × *s* identity matrix, ***D*** is the discrete diffusion operator ∇ ⋅ *D*∇ and ***A*** is an averaging matrix that assigns the density of the advected particle in space to the cell centers. This is derived via operator splitting where the advective step is performed using a particle-in-cell method and the backward Euler method is used for the diffusive step, see [REF] for more details. Notice, the density for any time step now depends only on the velocity ***v*** and initial density ***μ***_0_. Considering that the initial density is fixed by the given image ***μ***_0_ = ***μ***_*init*_, we have reduced the parameter space of our objective cost functional (1), *so that we only need find the optimal velocity.*

A straightforward discretization of the OMT energy functional 𝓡[*v*] (1) (as just noted we ignore the density term) is given as

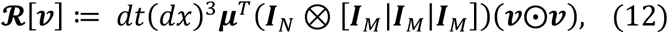

where ⨂ denotes the Kronecker product, [⋅ | ⋅] denotes block matrices and ⨀ denotes the Hadamard product. We seek the velocity *v* that characterizes the density ***μ*** which interpolates between the given initial and final densities, ***μ***_*init*_ and ***μ***_*final*_, seen to evolve according to the advection/diffusion equation (11) with minimal energy, as described by the discrete objection function **𝓡**[***v***] (12). Lastly, in anticipation of noise in the signal intensity, we do not attempt to match the final density exactly (3). Instead, we explicitly regard the final image as a noisy state of the tracer distribution

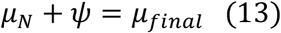

where *ψ* is independent, identically distributed Gaussian noise with covariance Ψ, treated as a free temporal endpoint of the partial differential equation. As such, the discrete regularized OMT problem is given as

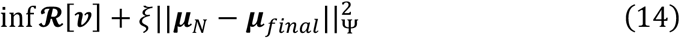

such that

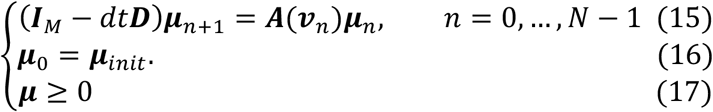

Here, the free-endpoint condition added to the objective function (14) inhibits the erroneous influence of noise on the velocity, inversely weighted by the covariance and the parameter ξ indicates the relative significance of fitting the data to minimizing the transport energy.

Using no diffusion or too small of a diffusion value yields implausible results (e.g. non-smooth velocity fields due to overcompensation by advection which therefore cannot reflect proper trajectories), but too large a value will have an accumulated effect that may end up over-smoothing the images over time (e.g. intensity gradients will be blurred yielding fictitious advective behavior attempting to reverse the spreading and subtler information will be lost). Multiple levels of diffusion were tested to determine values that afforded an appropriate balance. We note that for small changes in diffusion values (i.e. changes of the same order of magnitude), we found the model to be robust to the diffusivity. The aforementioned implausible behavior was only observed when the magnitude of the diffusivity was changed by an order of 2 or more, as expected. One would expect that increasing diffusion in a system of diffuse particles would allow for more random or unexpected behavior to occur. In this manner, diffusion allows for a natural description of stochastic particle trajectories thereby regularizing the flow. Accordingly, adding the diffusion term to the classical OMT constraint enables our model to account for more behavior than would be possible with advection alone. We therefore refer to our system of equations (14)–(17) as the ***free regularized OMT problem (frOMT), used interchangeably with rOMT)***, which turns out to be a quadratic optimization problem with linear constraints with respect to ***v***. The optimal velocity and interpolated densities are then found using a Gauss-Newton method.

### rOMT data analysis

DCE-MRI images were fed as inputs into our rOMT model in order to capture advective and diffusive transport in different compartments including CSF spaces, grey matter (GM), white matter (WM), brain regions (and PVS) along the MCA. Unless otherwise specified, all computation was carried out in MATLAB (Version 2016b). The rOMT procedure was run in parallel on all 14 whole brain datasets (8 WKY, 6 SHRSP) on the SeaWulf (Stony Brook University) CPU cluster (with a total of 4592 nodes and 128 gigabytes RAM per node) to model a 120 min time period of tracer transport captured across 21 images and took approximately 52 hours to execute. We note that fewer interpolated time steps could have been used to achieve a shorter runtime, but we felt a thorough investigation warranted the extended time. rOMT was performed on the 8 higher resolution MCA datasets (4 WKY and 4 SHRSP) in the same manner over a time period of 144 min captured across 25 images with a total runtime of approximately 66 hrs. More time was required due to the longer time period being considered and decreased signal-to-noise ratio but again, fewer intermediate steps could be used with minimal differences in output for future applications.

We used the Lagrangian OMT formulation (*vide supra*) to construct the pathlines for visualizing GS transport flows over the 120 min time period in one comprehensive figure, derived from the rOMT returned velocity field and interpolated images. In order to extract and visualize trajectories from 4D data in a meaningful way, the several steps were performed on each dataset. First, pathline start points were selected uniformly among voxels that showed at least 12% increase in signal as compared to baseline throughout the imaging acquisition interval. This was done in order to reduce the complexity of the visualization, as the trajectory of a parcel from a location with such low fluctuation may be attributed to noise or insignificant behavior. Pathlines were then parametrized by the Lagrangian coordinates (15) associated with each start point and were computed by integrating the augmented time-varying velocity field (13), as described above. Particle attributes such as speed, informing on the state of the particle along its trajectory, were simultaneously recorded and referred to as *speed-lines*. The pathlines were subsequently clustered by spatial proximity using the QuickBundles algorithm^66^ in Python (Version 3.5.4), and pathlines of small clusters deemed ‘insignificant’, as determined by a predefined threshold, were removed from consideration. ‘Significant’ pathlines were then converted back from spatial coordinates to the cell centered grid using a tri-linear averaging matrix and saved in nifty format for visualization in Amira. Runtime for the Lagrangian procedure is ~3 min per dataset on a standard laptop with a 2.2 GHz Intel Core i7 processor.

Time-varying particle attributes associated with the pathlines, such as speed and density, provide more detailed information regarding the particle dynamics. We selectively focus on the speed for the novel insight it provides. Specifically, speeds that makeup the aforementioned speedlines are computed as the magnitudes of the rOMT-derived transport vectors ascribed to the time and place of each point along the particle’s path and then visualized to analyze speed trajectories in different brain regions and across normotensive and hypertensive states.

For each rat, one Lagrangian rOMT image reflecting pathlines and one reflecting pathline speed (i.e. speedlines) at a voxel resolution of 0.3×0.3×0.3 for whole brain and 0.15×0.15×0.15 mm^3^ for hemispheric data were computed. For the whole brain data, total pathline volume and mean pathline speed were extracted from CSF, GM, WM and four predetermined brain regions (basal forebrain, superior colliculus, hippocampus and cerebellum). For MCA data, pathline volume and mean pathline speed were extracted in regions along the MCA segment on the base of the brain (a.k.a. MCA ‘root’) and along the left MCA.

### Immunohistochemistry

Animals were deeply anesthetized with ketamine/dexmedetomidine (150 mg kg^−1^/2 mg kg^−1^ i.p) and were transcardially perfused with phosphate buffered saline (PBS) followed by paraformaldehyde (PFA; 4% in PBS). The brains were extracted, post-fixed overnight in PFA at 4°C and transferred to PBS. All incubations were carried out at room temperature unless specified. Brains were sectioned coronally (50 μm) with a vibratome (Leica VT1000 S) and underwent heat-induced epitope retrieval in citrate buffer (10 mM citric acid, 0.05% Tween 20, pH 6.0) at 85°C for 30 min. The sections were then blocked and permeabilized for 1.5 h (1% BSA, 1% normal goat serum, and 0.3% Triton) and incubated for 2 h at 25°C with primary antibodies: rabbit anti-AQP4 (Novus, NBP1-87679, 1:500) and chicken anti-GFAP (Abcam, ab4674, 1:1000) in PBS, 0.05% Tween 20, and 1% BSA. Following this, sections were incubated for 2 h at 25°C with Alexa Fluor-Plus goat anti-rabbit 555 and Alexa Fluor goat anti-chicken 647 secondary antibodies (Invitrogen, 1:1000) in PBS with 0.05% Tween 20. The sections were then mounted on slides and cover slipped.

The MCA brains were also processed for GFAP, AQP4 and collagen IV. The PFA fixed brains were cryoprotected in 30% (w/v) sucrose in PBS, embedded in O.C.T. compound and snap-frozen at −80°C. Sections were cut coronally (20 μm) using a cryostat, mounted on Colorfrost / Plus slides and stored at −80°C. The sections were rehydrated in PBS, incubated with proteinase K for 15 min (0.2 mg/ml) and incubated in blocking buffer for 30 min (0.3% Triton in SuperBlock buffer). The sections were then incubated overnight at 4°C with primary antibodies: goat anti-collagen IV (Novus, NBP-126549,1:200), rabbit anti-GFAP (Dako, Z0334, 1:250), rabbit anti-AQP4 (Novus, NBP-1-87679) and goat anti-Iba-1 (Novus, NBP-100-1028, 1:250). The next day, sections were incubated with Alexa Fluor 594 or 488 secondary antibodies (Invitrogen, 1:1000). The sections were then cover slipped and sealed. Images were captured using a Keyence fluorescence microscope BZ-X700 system.

### Quantification of AQP4 polarization

Images of AQP4 immunofluorescence were captured on a Keyence BZ-X700 automated fluorescence microscope using 4-20 × air objectives as *z*-stack tile-scans with 3 μm *z*-steps in maximum contrast projections. Quantification of AQP4 polarization was done in the ventral hippocampus and occipital cortex using ImageJ software (Image J 1.52i). The AQP4 digitized images were imported into ImageJ, scaled and converted to optical density images. In order to unbias the analysis, a grid (150 numbered boxes) was overlaid over the ventral hippocampus (each box area = 228 × 228 pixels) and occipital cortex (each box area = 282 × 282 pixels). 15 boxes from hippocampal or cortical sections were randomly selected for a total of ~30 small vessels and ~30 capillaries for WKY rats and the same number of vessels in SHRSP rats. Representative line segments (300 μm for small vessels) and 100 μm for capillaries were used to extract immuno-intensity across the micro-vessels selected in each grid-box. The polarization index for each vessel was calculated (peak-to-baseline level).

### Quantification of GFAP and AQP4 mRNA by real-time quantitative RT-PCR (qPCR)

Astrogliosis and AQP4 water channel expression were evaluated in the brains of WKY and SHRSP rats suing real-time quantitative RT-PCR (qPCR). Specifically, for this purpose, mRNA levels of the two targets (GFAP and AQP4) by qPCR was performed with the QuantStudio 3 Real-Time PCR system (Applied Biosystems). Brain tissues of interest (WKY, N=8; SHRSP, N=8) were quickly dissected, snap frozen in liquid nitrogen and stored at −80 °C. Total RNA extraction was carried out using the PARIS kit (Invitrogen) following brief homogenization using the TissueRuptor II (Qiagen). Total RNA concentrations were measured using the SpectraMax i3x Multi-Mode microplate reader (Molecular Devices). Samples were then treated with ezDNase enzyme (Invitrogen) to degrade any genomic DNA and subsequently reverse transcribed to cDNA using the Superscipt IV VILO kit (Invitrogen). Quantification of target mRNA was performed using TaqMan gene expression assays (ThermoFisher Scientific). The PCR cycle parameters were set according to the recommended guidelines for the TaqMan Fast Advanced Master Mix (Applied Biosystems), which was used to run the qPCR reactions (95 °C for 2 min [polymerase activation] followed by 40 cycles of 95 °C for 1 s [denture] and 60 °C for 20 s [anneal/extend]). TaqMan primer/probe sets for AQP4 (Rn00563196_m1) and GFAP (Rn00566603_m1) were used. mRNA target signals were normalized to the housekeeping gene RPLP0 (Rn00821065_g1). Relative fold change values were calculated using the 2^−ΔΔCt^ method^67^. The mRNA target signals were firstly normalized to the reference housekeeping gene RPLP0. Following this, relative fold change values were calculated using the 2^−ΔΔCt^ method by normalizing the data to the cerebellum, which was used a reference region.

### Statistical information

Group sizes for the whole brain glymphatics studies were estimated based on previous studies^37, 59^. For peak magnitude point comparisons, group sample sizes of 6 (WKY) and 6 (SHRSP) achieve 80% power to detect a Cohen’s effect size of 2.0, and with a significance level (α) of 0.025 using a two-sided two-sample unequal-variance t-test. All data are presented as mean ± SD. If data showed normal distribution, a paired or unpaired Student’s t-test was used to compare groups. When data lacked normal distribution, groups were compared using non-parametric testing (Mann-Whitney test for unpaired comparison, Wilcoxon signed rank test for paired comparisons). For within-group regional differences Friedman’s ANOVA was used. A at a 0.05 level of significance and stated in the figure legends. Blinding for data analysis was done where possible. All statistical analyses were performed using XLSTAT statistical and data analysis solution (Addinsoft (2019). XLSTAT statistical and data analysis solution. Boston, USA. https://www.xlstat.com) or GraphPad Prism 7 (GraphPad Software).

## Acknowledgements

NIH R01AG048769, RF1 AG053991, R01AG057705 and Leducq Foundation (16/CVD/05)

## Supplementary Information

**Supplementary Fig. 1:**
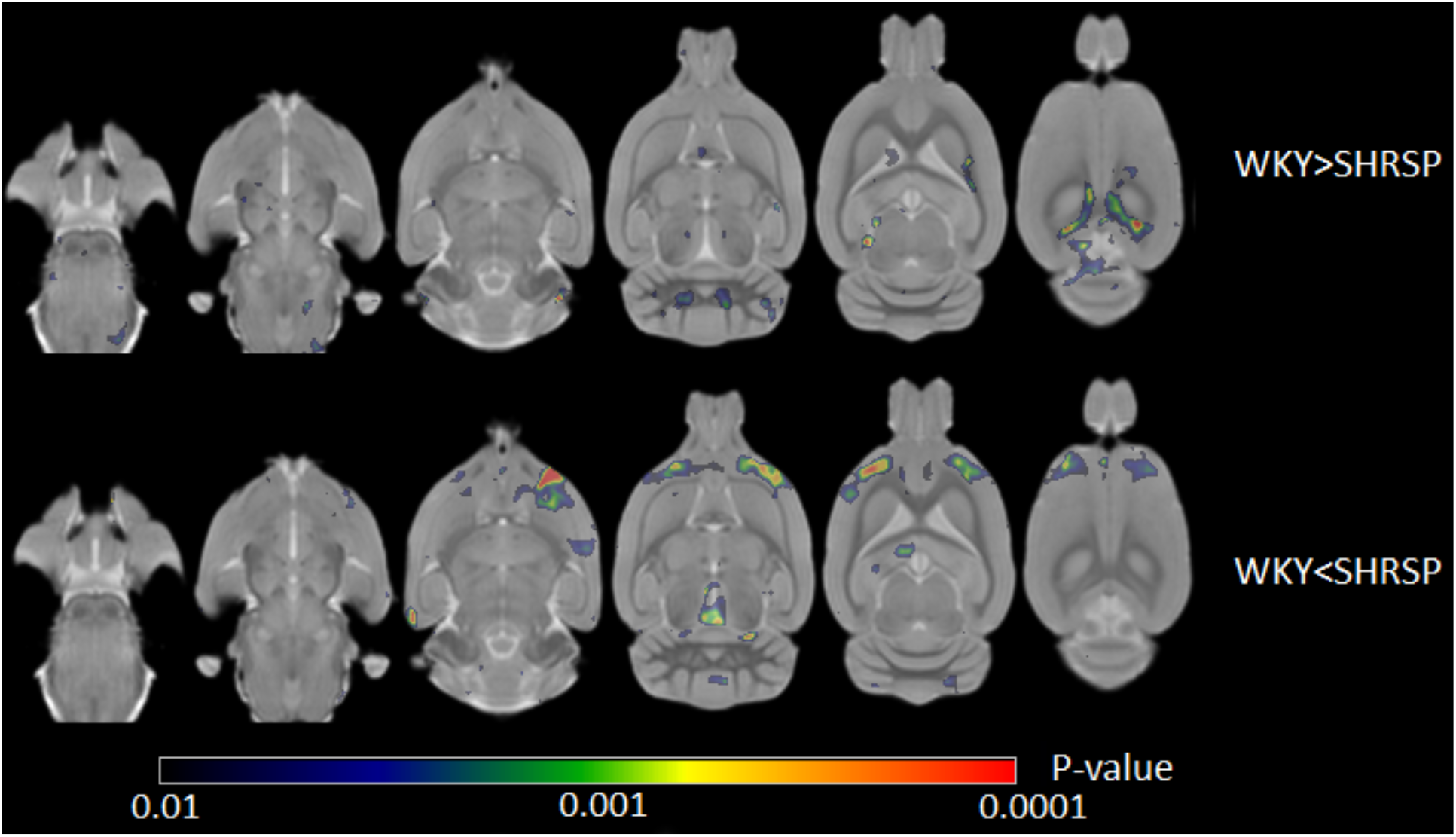
Morphometric brain analysis of WKY and SHRSP rats. Voxel wise deformation-based morphometry results displayed as a color-coded p-value map overlaid on spatially normalized, population averaged proton-density weighted anatomical MRI brain images of WKY (n=6) and SHRSP (n=6) rats. The top series of images shows regions that are larger in WKY compared to SHRSP rats, including areas associated with the white matter of cerebellum and the splenium of corpus callosum. The bottom series of images highlights areas that are significantly larger in SHRSP than WKY rats, including frontal cortical regions, the periaqueductal grey, and the aqueduct.

**Supplementary Fig. 2:**
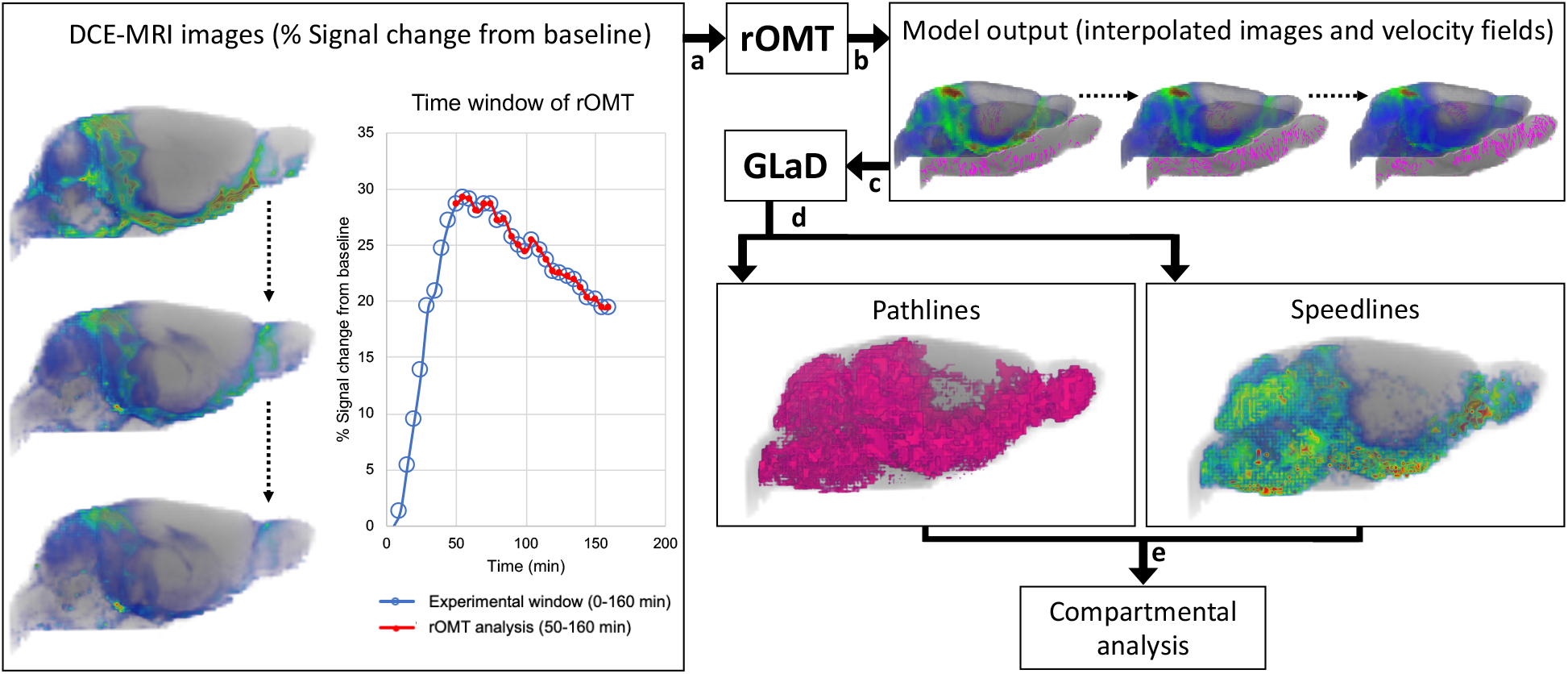
Regularized optimal mass transport (rOMT) and *La*grangian representation of *Gl*ymphatics *D*ynamics (GLaD) pipeline for visualizing transport flows over a predetermined time interval. **(a)** DCE-MRIs over a 120 min experimental window (50-160 min) were fed as inputs into the rOMT model (**b**) The model returns the interpolated density images and velocity fields which are (**c**) subsequently processed by our GLaD visualization framework. (**d**) Advective and diffusive GS transport is expressed by the pathlines captured in the Lagrangian analysis with corresponding dynamic speedlines. Binary pathlines were used to extract flow volume and speedlines were used to assess spatial distribution of speed trajectories in whole brain as well as to derive mean speed values. (**e**) Thereafter, compartmental analysis is carried out based on tissue masks derived from the morphometric analysis.

**Supplementary Fig. 3:**
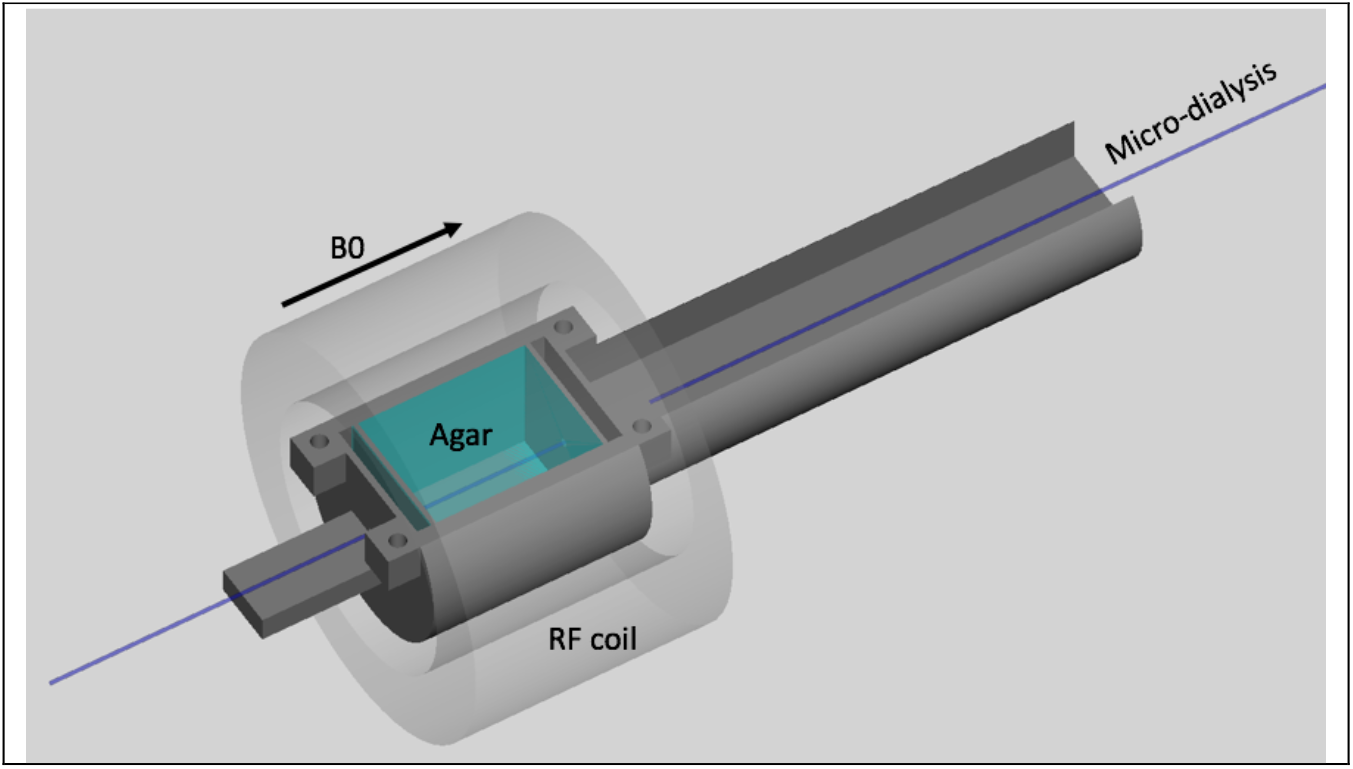
Diffusion phantom with microdialysis tube. We validated that the rOMT algorithm captures Fick’s law using a custom built ‘diffusion’ phantom with a microdialysis membrane. A linear designed microdialysis probe (outer diameter 260μm, molecular weight cut off 55kDa, CMR 31) was inserted through the center of a 3D printed, MR compatible holder containing 0.5% agar (20×20×30 mm). The phantom was placed in the 9.4T MRI scanner equipped with 40 mm ID RF transmit/receive coil and the microdialysis membrane was perfused for 43 min (108 μl) with Gd-DOTA (12 mM) at 2.5 μl/min using a micro-infusion pump after which Gd-DOTA solutes moved into the agar medium driven by diffusion. Pre- and post-contrast DCE-MRI were acquired using the same parameters as for the *in vivo* study. A total of 140 frames were collected at an interval of 2.5 min per frame under constant temperature (21-22°C).

**Supplementary Fig. 4.**
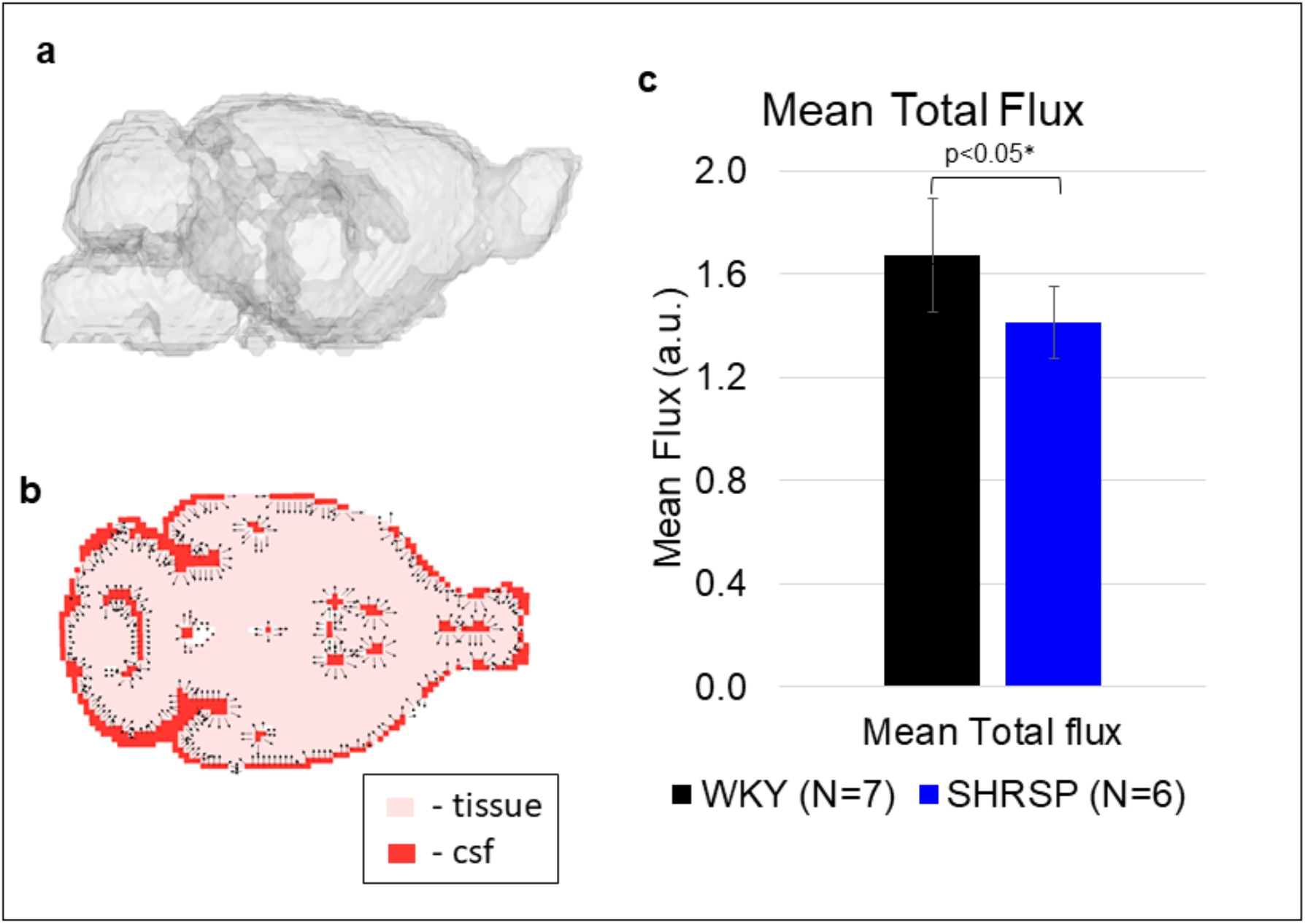
rOMT relation to compartmental models. **(a)** 3D rendering of boundary *B* of tissue compartment *C*. (**b)** Representative slice of the whole brain showing the CSF compartment in red and the tissue compartment (gray matter and white matter) in pink. Normal vectors (of unit length) pointing from the CSF compartment into the tissue compartment are shown along the CSF-tissue boundary as black arrows. (**c)** Mean (advective + diffusive) flux that moved from CSF into the tissue compartment is significantly higher in normotensive (WKY) than hypertensive (SHRSP) cases.

**Supplementary Fig. 5:**
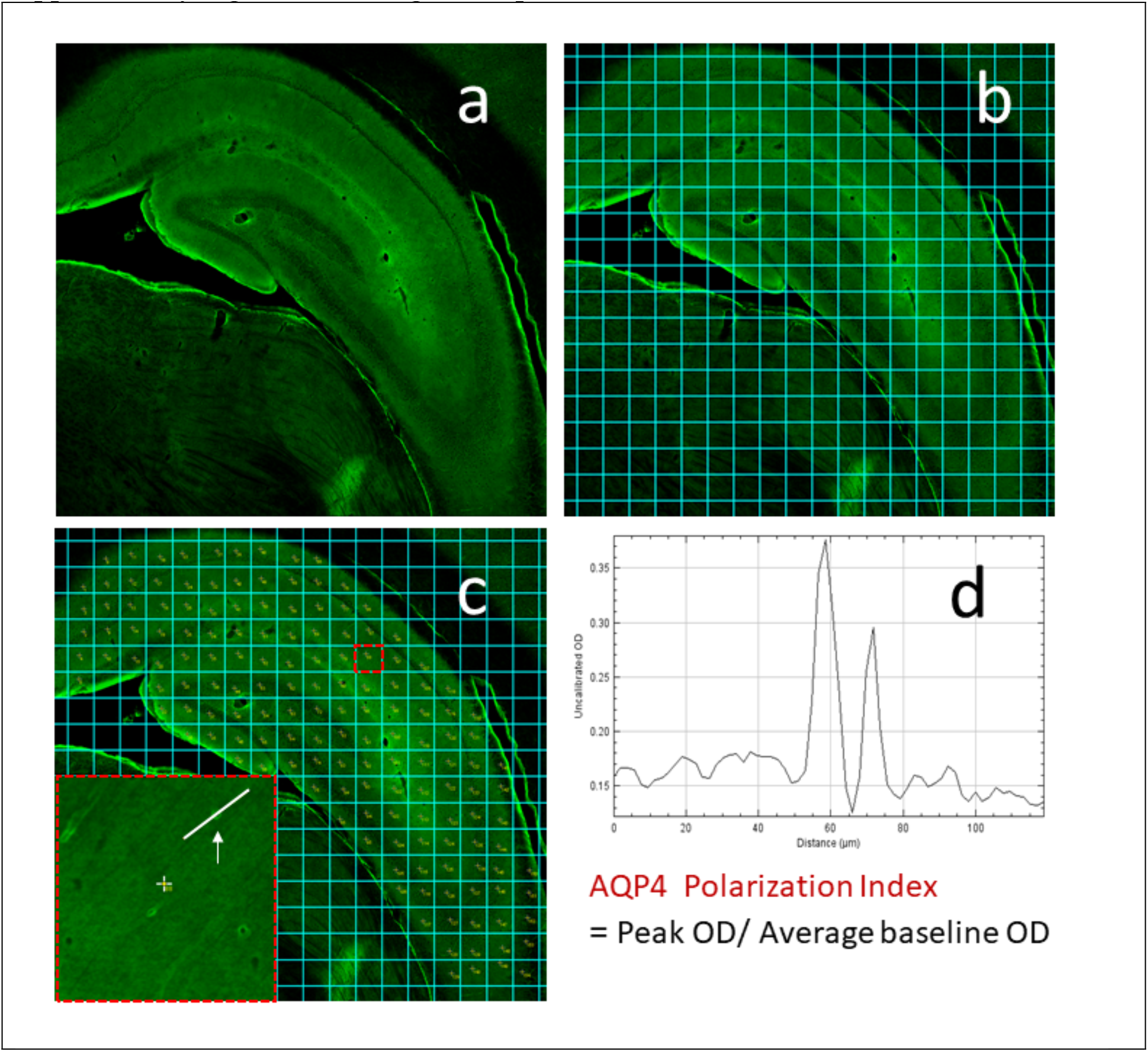
(**a)** AQP4 immunofluorescence at the level of ventral hippocampus from a WKY rat. **(b)** For quantification of AQP4 polarization, a uniform, 154 box grid was overlaid using ImageJ (Image J 1.52i). **(c)** 10 and 15 random numbers were selected from the 154-box grid for measurement of small vessels and capillaries, respectively. The arrow points to the linear ROIs crossing the lumen of the small vessel of interest in the grid of interest. **(d)** The optical density profile from the ROI in (c) is displayed along with the calculated polarization index. Specifically, for polarity calculation, the peak intensity of the vascular end-feet was divided by the average of the baseline.

**Supplementary Fig. 6:**
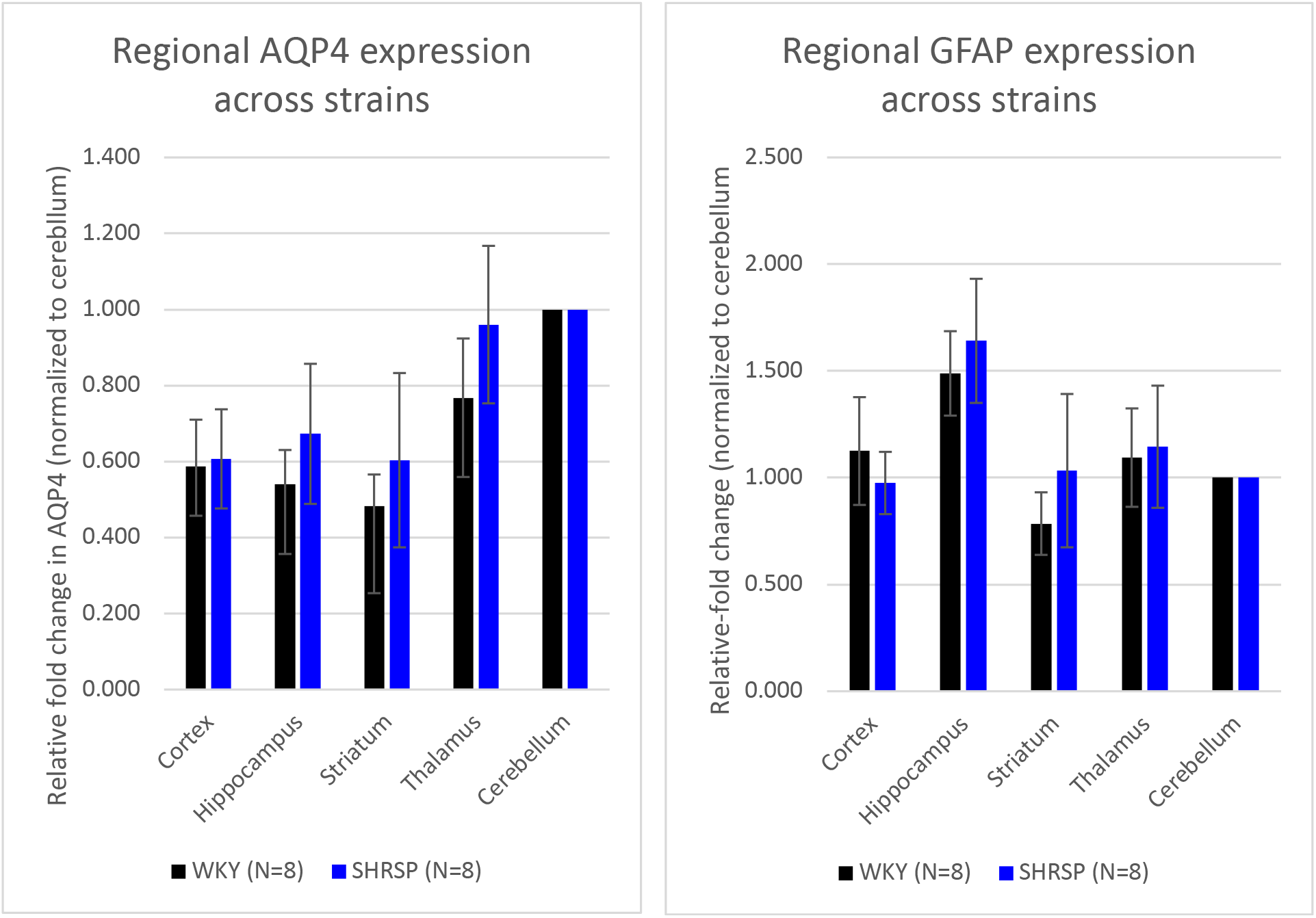
qPCR analysis of GFAP and AQP4 expression in WKY and SHRSP rats. qPCR analysis of AQP4 (left) and GFAP (right) mRNA expression in WKY (n=8) and SHRSP (n=8) rats assessed in cortex, hippocampus, striatum and thalamus. Relative fold change values were calculated by normalizing to the cerebellum. Data are expressed as mean relative fold change values ± standard deviation. No significant differences were found between regional AQP4 and GFAP mRNA expression across strains. All statistical analyses were performed using Addinsoft (2019). XLSTAT statistical and data analysis solution. Boston, USA. https://www.xlstat.com.

**Supplementary Fig. 7:**
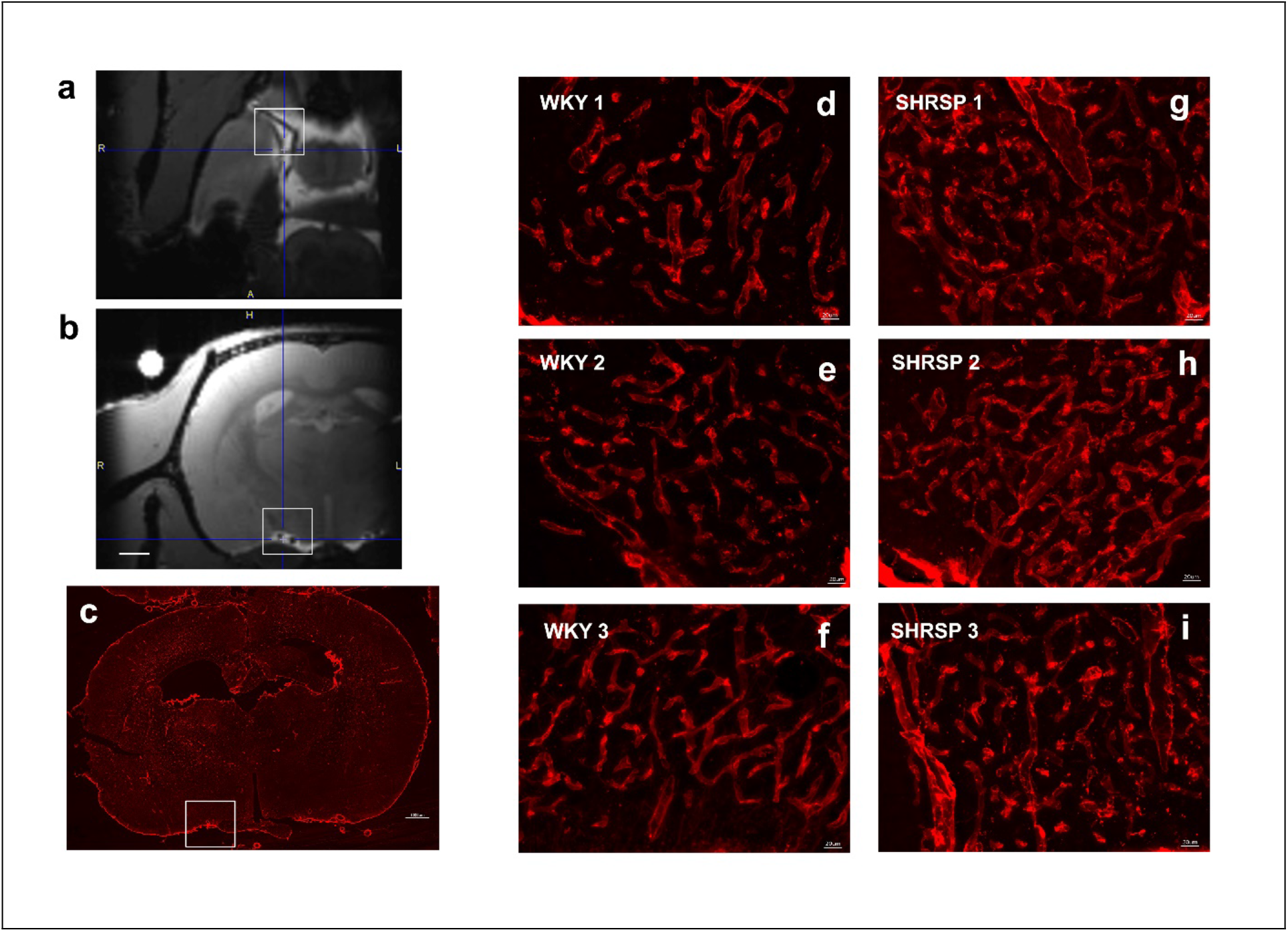
Exploration of the microvasculature in WKY and SHRSP at the level of circle of Willis. **(a, b)** Contrast enhanced MRIs from a WKY rat at the level of the circle of Willis near the take off of the middle cerebral artery (MCA). Contrast in the perivascular space along the circle of Willis and middle cerebral artery is evidenced as a high intensity signal along the vasculature (black signal). **(c)** Collagen IV immunofluorescence of a WKY rat. Insert shows the area of interest near the large vessels at the ventral surface of the brain. (**d-f)** Representative confocal montages of collagen IV labeling from three different WKY rats at the site highlighted by the white box in (c). Smaller networks of capillaries (7-10 μm in diameter) were observed at the base of the brain surrounding penetrating arteries and appear interweaved into a small network. **(g-i)** In corresponding images from the SHRSP rats, collagen IV labeling showed the capillary networks (a.k.a. rete) were distributed over slightly larger areas around penetrating arteries. Further, the capillary networks appeared denser and the individual vessels were larger (10-20 μm in diameter). We speculate that the microvascular network remodeling might have affected the PVS transport from the circle of Willis in SHRSP compared to WKY rats.

**Supplemental Movie 1. Regularized OMT model (rOMT) captures diffusive agar phantom behavior.** rOMT-returned interpolated images (right) are consistent with DCE-MRI phantom data (left). rOMT successfully models diffusive transport from regions of higher intensity to regions of lower intensity in accordance with Fick’s law, shown by diffusive flux vectors in green. See Movie1.mov.

**Supplementary Movie 2:** Contrast-enhanced dynamic MRI from normal WKY rat showing signal changes along the middle cerebral artery over ≈3hr CSF Gd-DOTA circulation. See Movie2.mpg.

**Supplementary Table 1:** Physiological data and anesthetics.

| Experiment | Parameter | Animals anesthetized? | WKY (N=11) | SHRSP (N=11) | p-value |
| --- | --- | --- | --- | --- | --- |
| **Non-invasive Blood pressure** | Systolic blood pressure (mmHg) | No | 116 ± 14 | 150 ± 23 | **0.0004** |
| | Diastolic blood pressure (mmHg) | No | 85 ± 11 | 98 ± 21 | 0.151 |
| | Pulse pressure (mmHg) | No | 32 ± 9 | 51 ± 13 | **0.001** |
| Experiment | Parameter | Animals anesthetized? | WKY (N=6) | SHRSP (N=5) | p-value |
| **Invasive blood pressure** | Mean arterial blood pressure (mmHg) | Yes | 76 ± 14 | 123 ± 25 | **0.004** |
| Experiment | Parameter | Animals anesthetized? | WKY (N=11) | SHRSP (N=10) | p-value |
| **Glymphatics Experiments** | Mean heart rate (bpm) | Yes | 221 ± 20 | 235 ± 14 | 0.049 |
| | Mean respiratory rate (breath/min) | Yes | 45 ± 6 | 46 ± 7 | 0.650 |
| | Mean infusion rate of dexmedetomidine* (ml/hr) | N/A | 0.10 ± 0.03 | 0.10 ± 0.03 | 0.748 |
| | Mean inspiratory isoflurane concentration (%) | N/A | 1.1 ± 0.2 | 1.0 ± 0.2 | 0.932 |

Data are mean ± SD. \*Concentration of dexmedetomidine used for infusion: 0.025mg/ml.

## References

1. Iliff JJ, et al. A paravascular pathway facilitates CSF flow through the brain parenchyma and the clearance of interstitial solutes, including amyloid beta. Science translational medicine 4, 147ra111 (2012).

2. Nedergaard M. Neuroscience. Garbage truck of the brain. Science 340, 1529–1530 (2013).

3. Iliff JJ, et al. Impairment of glymphatic pathway function promotes tau pathology after traumatic brain injury. The Journal of neuroscience : the official journal of the Society for Neuroscience 34, 16180–16193 (2014).

4. Benveniste H, Liu X, Koundal S, Sanggaard S, Lee H, Wardlaw J. The Glymphatic System and Waste Clearance with Brain Aging: A Review. Gerontology 65, 106–119 (2019).

5. Brown R, et al. Understanding the role of the perivascular space in cerebral small vessel disease. Cardiovasc Res, (2018).

6. Zeppenfeld DM, et al. Association of Perivascular Localization of Aquaporin-4 With Cognition and Alzheimer Disease in Aging Brains. JAMA Neurol 74, 91–99 (2017).

7. Eide PK, Ringstad G. Delayed clearance of cerebrospinal fluid tracer from entorhinal cortex in idiopathic normal pressure hydrocephalus: A glymphatic magnetic resonance imaging study. J Cereb Blood Flow Metab, 271678X18760974 (2018).

8. Iliff JJ, et al. Brain-wide pathway for waste clearance captured by contrast-enhanced MRI. J Clin Invest 123, 1299–1309 (2013).

9. Cserr HF. Role of secretion and bulk flow of brain interstitial fluid in brain volume regulation. Ann N Y Acad Sci 529, 9–20 (1988).

10. Cserr HF, Depasquale M, Patlak CS, Pullen RG. Convection of cerebral interstitial fluid and its role in brain volume regulation. Ann N Y Acad Sci 481, 123–134 (1986).

11. Weed LH. Studies on Cerebro-Spinal Fluid. No. III : The pathways of escape from the Subarachnoid Spaces with particular reference to the Arachnoid Villi. J Med Res 31, 51–91 (1914).

12. Smith AJ, Verkman AS. The “glymphatic” mechanism for solute clearance in Alzheimer’s disease: game changer or unproven speculation? FASEB J 32, 543–551 (2018).

13. Smith AJ, Yao X, Dix JA, Jin BJ, Verkman AS. Test of the ‘glymphatic’ hypothesis demonstrates diffusive and aquaporin-4-independent solute transport in rodent brain parenchyma. Elife 6, (2017).

14. Jin BJ, Smith AJ, Verkman AS. Spatial model of convective solute transport in brain extracellular space does not support a “glymphatic” mechanism. J Gen Physiol 148, 489–501 (2016).

15. Abbott NJ, Pizzo ME, Preston JE, Janigro D, Thorne RG. The role of brain barriers in fluid movement in the CNS: is there a ‘glymphatic’ system? Acta Neuropathol 135, 387–407 (2018).

16. Smith AJ, Verkman AS. CrossTalk opposing view: Going against the flow: interstitial solute transport in brain is diffusive and aquaporin-4 independent. J Physiol, (2019).

17. Smith AJ, Verkman AS. Rebuttal from Alex J. Smith and Alan S. Verkman. J Physiol, (2019).

18. Simon M, Iliff J. Rebuttal from Matthew Simon and Jeffrey Iliff. J Physiol, (2019).

19. Iliff JJ, et al. Cerebral arterial pulsation drives paravascular CSF-interstitial fluid exchange in the murine brain. The Journal of neuroscience : the official journal of the Society for Neuroscience 33, 18190–18199 (2013).

20. Rennels ML, Gregory TF, Blaumanis OR, Fujimoto K, Grady PA. Evidence for a ‘paravascular’ fluid circulation in the mammalian central nervous system, provided by the rapid distribution of tracer protein throughout the brain from the subarachnoid space. Brain Res 326, 47–63 (1985).

21. Mestre H, et al. Flow of cerebrospinal fluid is driven by arterial pulsations and is reduced in hypertension. Nat Commun 9, 4878 (2018).

22. Abbott NJ. Evidence for bulk flow of brain interstitial fluid: significance for physiology and pathology. Neurochem Int 45, 545–552 (2004).

23. Holter KE, et al. Interstitial solute transport in 3D reconstructed neuropil occurs by diffusion rather than bulk flow. Proc Natl Acad Sci U S A 114, 9894–9899 (2017).

24. Asgari M, de Zelicourt D, Kurtcuoglu V. Glymphatic solute transport does not require bulk flow. Sci Rep 6, 38635 (2016).

25. Gupta S, Soellinger M, Boesiger P, Poulikakos D, Kurtcuoglu V. Three-dimensional computational modeling of subject-specific cerebrospinal fluid flow in the subarachnoid space. J Biomech Eng 131, 021010 (2009).

26. Rey J, Sarntinoranont M. Pulsatile flow drivers in brain parenchyma and perivascular spaces: a resistance network model study. Fluids Barriers CNS 15, 20 (2018).

27. Keith Sharp M, Carare RO, Martin BA. Dispersion in porous media in oscillatory flow between flat plates: applications to intrathecal, periarterial and paraarterial solute transport in the central nervous system. Fluids Barriers CNS 16, 13 (2019).

28. Rachev ST, Rüschendorf L. Mass Transportation Problems. Springer (1998).

29. Ratner V, et al. Cerebrospinal and interstitial fluid transport via the glymphatic pathway modeled by optimal mass transport. NeuroImage 152, 530–537 (2017).

30. Yongxin C, Eldad H, Kaoru Y, T. Gt, Tannenbaum A. An Efficient Algorithm for Matrix-Valued and Vector-Valued Optimal Mass Transport. J Scientific Computing 77, 79–100 (2018).

31. Brenier Y, et al. Reconstruction of the early Universe as a convex optimization problem. Mon Not R Astron Soc 346, 501–524 (2003).

32. Kolouri S, Park SR, Thorpe M, Slepcev D, Rohde GK. Optimal Mass Transport Signal processing and machine-learning applications. Ieee Signal Proc Mag 34, 43–59 (2017).

33. Villani C. Optimal Transport: Old and New. Grundlehr Math Wiss 338, 1–973 (2009).

34. Benamou JD, Brenier Y. A computational fluid mechanics solution to the Monge-Kantorovic mass transfer problem. Numirische Mathematik 84, 375–393 (2000).

35. Peng W, et al. Suppression of glymphatic fluid transport in a mouse model of Alzheimer’s disease. Neurobiology of disease 93, 215–225 (2016).

36. Kress BT, et al. Impairment of paravascular clearance pathways in the aging brain. Annals of neurology 76, 845–861 (2014).

37. Nygaard Mortensen K, et al. Impaired Glymphatic Transport in Spontaneously Hypertensive Rats. The Journal of neuroscience : the official journal of the Society for Neuroscience, (2019).

38. Jiang Q, et al. Impairment of the glymphatic system after diabetes. J Cereb Blood Flow Metab 37, 1326–1337 (2017).

39. Lee H, et al. Quantitative Gd-DOTA uptake from cerebrospinal fluid into rat brain using 3D VFA-SPGR at 9.4T. Magn Reson Med 79, 1568–1578 (2018).

40. Bedussi B, et al. Enhanced interstitial fluid drainage in the hippocampus of spontaneously hypertensive rats. Sci Rep 7, 744 (2017).

41. Koundal S, et al. Brain Morphometry and Longitudinal Relaxation Time of Spontaneously Hypertensive Rats (SHRs) in Early and Intermediate Stages of Hypertension Investigated by 3D VFA-SPGR MRI. Neuroscience 404, 14–26 (2019).

42. Ritter S, Dinh TT. Progressive postnatal dilation of brain ventricles in spontaneously hypertensive rats. Brain Res 370, 327–332 (1986).

43. Ghosh D, Ghosh PP, Gambhir S, Kohli A. Normal pressure hydrocephalus role of radionuclide cisternography. Neurol India 45, 231–239 (1997).

44. Benveniste H, Huttemeier PC. Microdialysis--theory and application. Prog Neurobiol 35, 195–215 (1990).

45. Nicholson C, Sykova E. Extracellular space structure revealed by diffusion analysis. Trends in neurosciences 21, 207–215 (1998).

46. Xie L, et al. Sleep drives metabolite clearance from the adult brain. Science 342, 373–377 (2013).

47. Benveniste H, et al. Anesthesia with Dexmedetomidine and Low-dose Isoflurane Increases Solute Transport via the Glymphatic Pathway in Rat Brain When Compared with High-dose Isoflurane. Anesthesiology, (2017).

48. Mestre H, et al. Aquaporin-4-dependent glymphatic solute transport in the rodent brain. Elife 7, (2018).

49. Monge G. Mémoire sur la théorie des déblais et des remblais. . In: Histoire de l’Académie Royale des Sciences de Paris, avec les Mémoires de Mathématique et de Physique pour la même année (ed^(eds). Paris: De I’Imprimerie Royale (1871).

50. Lee H, et al. Quantitative Gd-DOTA uptake from cerebrospinal fluid into rat brain using 3D VFA-SPGR at 9.4T. Magn Reson Med In Press, (2017).

51. White FM. Viscous fluid flow, 3rd edn. McGraw-Hill Higher Education (2006).

52. White FM, Corfield I. Viscous fluid flow. McGraw-Hill (2006).

53. Lee H, et al. The Effect of Body Posture on Brain Glymphatic Transport. The Journal of neuroscience : the official journal of the Society for Neuroscience 35, 11034–11044 (2015).

54. Davoodi-Bojd E, et al. Modeling glymphatic system of the brain using MRI. NeuroImage 188, 616–627 (2019).

55. Horn B. Robot vision, MIT Press edn. MIT Press; McGraw-Hill (1986).

56. Wardlaw JM, Smith C, Dichgans M. Mechanisms of sporadic cerebral small vessel disease: insights from neuroimaging. The Lancet Neurology 12, 483–497 (2013).

57. Pedder H, Vesterinen HM, Macleod MR, Wardlaw JM. Systematic review and meta-analysis of interventions tested in animal models of lacunar stroke. Stroke 45, 563–570 (2014).

58. Chen Y, Georgiou TT, Tannenbaum A. Interpolation of matrices and matrix-valued densities: The unbalanced case. European Journal of Applied Mathematics 30, 1–23 (2019).

59. Benveniste H, et al. Anesthesia with Dexmedetomidine and Low-dose Isoflurane Increases Solute Transport via the Glymphatic Pathway in Rat Brain When Compared with High-dose Isoflurane. Anesthesiology 127, 976–988 (2017).

60. Kjonigsen LJ, Lillehaug S, Bjaalie JG, Witter MP, Leergaard TB. Waxholm Space atlas of the rat brain hippocampal region: three-dimensional delineations based on magnetic resonance and diffusion tensor imaging. NeuroImage 108, 441–449 (2015).

61. Ashburner J. A fast diffeomorphic image registration algorithm. NeuroImage 38, 95–113 (2007).

62. Ashburner J, Friston KJ. Unified segmentation. NeuroImage 26, 839–851 (2005).

63. Chen Y, Georgious TT, Payon M. On the relation between optimal transport and Schrödinger bridges: A stochastic control viewpoint. Journal of Optimization Theory and Applications 169, 671–691 (2016).

64. Tofts PS, et al. Estimating kinetic parameters from dynamic contrast-enhanced T(1)- weighted MRI of a diffusable tracer: standardized quantities and symbols. J Magn Reson Imaging 10, 223–232 (1999).

65. Steklova K, Haber E. Joint hydrogeophysical inversion: state estimation for seawater intrusion models in 3D. Computat Geosci 21, 75–94 (2017).

66. Garyfallidis E, Brett M, Correia MM, Williams GB, Nimmo-Smith I. QuickBundles, a Method for Tractography Simplification. Front Neurosci 6, 175 (2012).

67. Livak KJ, Schmittgen TD. Analysis of relative gene expression data using real-time quantitative PCR and the 2(T)(-Delta Delta C) method. Methods 25, 402–408 (2001).

